# Engineering a Multilayer Microfluidic Airway-On-A-Chip with Tunable GelMA Hydrogel for Physiologically Relevant Aerosol Exposure Studies

**DOI:** 10.1101/2025.11.06.687056

**Authors:** Tanya J. Bennet, Avineet Randhawa, Tara M. Caffrey, Tanya Solomon, Eric Lyall, Ryan D. Huff, Carley Schwartz, Yu Xi, Christopher Carlsten, Karen C. Cheung

## Abstract

Climate change-driven increases in forest fires pose a major global health risk due to exposure to smoke containing hazardous gases and fine particulates, emphasizing the need for physiologically relevant *in vitro* airway models for studying smoke-induced responses. Microfluidic lung-on-a-chip technologies provide a strong foundation for *in vitro* airway modeling and ongoing developments are expanding their ability to incorporate multicellular organization, extracellular matrix complexity, and physiologically relevant exposure methods. This work presents the optimization and integration of a photopolymerizable gelatin methacrylate (GelMA)-based hydrogel into a microfluidic airway-on-a-chip that models the human small conducting airways and supports controlled aerosol exposure to wood smoke. The GelMA hydrogel was optimized to support fibroblast encapsulation, endothelial, and epithelial adhesion and robust mechanical stability. The device combines the hydrogel with a compartmentalized microchannel layout, and sacrificial molding to create a 3D organotypic airway culture featuring a multilayer architecture, 3D stromal matrix, and a perfusable vasculature-like lumen. Coupling the platform with a custom aerosol exposure system enables precise, biomimetic exposure to whole wood smoke. Proof-of-concept studies using transforming growth factor beta1 (TGF-β1) and whole wood smoke elicited expected inflammatory and fibrotic responses, validating the platform’s physiological relevance for inhalation studies and investigating smoke-induced airway remodeling and inflammation.

## Introduction

Forest fires are a major source of air pollution and pose an increasing global health threat as climate change drives more frequent and severe events. Smoke from these fires contain harmful gases, chemicals, and fine particulate matter (PM2.5) that can penetrate deep into the lungs causing cell damage, inflammation, and impaired lung function^1,2^. Prolonged and repeated exposure can trigger chronic inflammation and extracellular matrix (ECM) remodeling, contributing to progressive and structural changes and diseases such as chronic obstructive pulmonary disease (COPD)^3,4^. Individuals with pre-existing respiratory conditions are especially vulnerable, experiencing worsened symptoms, hospitalization, and accelerated disease progression^3,4^. As climate change is expected to increase the number, size and duration of forest fires, associated health risks and economic burdens on the public health system are expected to rise, emphasizing the need to better understand mechanisms of wood smoke-induced cell damage and inflammation^5–9^.

Studying the health impacts of wildfire smoke is challenging due to the complexity of both the human airways and the resulting smoke and exposure dynamics. Cells are strongly influenced by their environment, especially in relation to dimensionality, physiological flows, and ECM cues yet most *in vitro* models are often oversimplified, failing to sufficiently replicate airway architecture, microenvironment, and functions, especially in the context of smoke-induced responses. Consequently, these models are limited in their ability to address important questions related to exposure. To advance understanding of smoke-related respiratory risks, more sophisticated *in vitro* models, capable of incorporating physiologically relevant aerosol-based inhalation exposures are needed.

Microfluidic lung-on-a-chip (LOC) technologies have emerged as promising tools for enhanced modeling as they can replicate native tissue structures, physiological airflow, and exposure to environmental pollutants in a biomimetic manner^10–15^. Since the lung consists of several distinct microenvironments, LOC models can be further classified based on their region of focus; alveoli, or airway. As the alveoli are the primary site of gas exchange, current LOC models favor this region over the airways often incorporating 2D monolayers on a basement membrane surrogate to recapitulate the epithelial-endothelial interface of the alveoli^16–25^. Airway LOC or Airway-on-a-chip (AOC) models differ in their anatomical focus, aiming to replicate cell architecture and interactions specific to the conducting airways^26–39^.

Fibroblasts play a critical role in airway homeostasis and remodeling, making their inclusion essential for modelling cell-ECM interactions and early responses to inhaled particulates. *Benam et al.*^40^, demonstrated an AOC model containing a co-culture of airway epithelial cells and endothelial cells for aerosolized cigarette smoke exposure. While their model demonstrates the benefits of microfluidic AOC models for inhalation studies, it lacks a 3D interstitial ECM and associated stromal cells, limiting its ability to capture cell-ECM interactions and suitability for studying airway remodeling and fibrosis. Similarly, membrane-separated models with 2D planar endothelial monolayers do not sufficiently mimic the 3D microvasculature geometry and surrounding ECM architecture that is critical for tissue function. Vascularization and ECM components are key features needed for accurately modeling fibrosis, a known smoke-related pathology (reviewed in^41^). Patterning hollow lumens within 3D hydrogels provides the opportunity to create physiologically relevant microvasculature models; however, few AOC models have utilized this approach^31^. Combining hydrogels with microfluidics also provides the basis for creating *in vivo*-like 3D microenvironments such as the air-liquid interface (ALI) in addition to replicating physiological flows and mechanical cues. Despite the advances in AOC models for studying exposures to airborne particulates or smoke^15^, many remain oversimplified, relying on liquid smoke extracts, or lacking dynamic airflow or realistic ECM environments. To harness the potential of LOC models, especially in the context of inhalation-related toxicity to environmental pollutants and associated fibrotic responses, it it’s essential to build 3D multicellular models capable of capturing both dynamic and complex cell-cell and cell-ECM interactions^41–43^.

Here, we report a novel microfluidic model of the small conducting airway, integrating a tunable gelatin methacrylate (GelMA) hydrogel compatible with whole smoke exposure. Hydrogel parameters were optimized for integration into the microfluidic device by modifying polymer and photoinitiator concentration, and sacrificial molding enabled the creation of an *in vivo*-like vasculature within the photopolymerizable hydrogel expanding the model’s physiological relevance and ability to capture cell-cell and cell-ECM interactions. To highlight its potential for inhalation research, the AOC was treated with transforming growth factor beta 1 (TGF-β1), a cytokine elevated in the airways during smoke exposure, and whole wood smoke.

## Results

### Hydrogel Selection and Optimization for AOC Integration

#### Hydrogel Selection: Rationale for Using Gelatin Methacrylate (GelMA)

GelMA, a semi-synthetic hydrogel derived from hydrolyzed collagen, was selected for the AOC platform for its biocompatibility, optical clarity, tunable mechanical properties and photopolymerization capability^44,45^. Methacrylation permits photopolymerization and the formation of covalent bonds, allowing it to be tunable for integration and long-term stability, while retaining gelatin’s cell-responsive features, including integrin binding sites (arginylglycylaspartic acid (RGD) peptide motif) and matrix metalloproteinase (MMP)-sensitive sites, providing an environment that supports cell adhesion, encapsulation, and remodeling^45,46^. GelMA hydrogels can be optimized for specific applications by adjusting polymer concentration, degree of functionalization, photoinitiator type and concentration, and light exposure conditions (wavelength and time)^47^. Since the airway ECM provides physiological support, biochemical signaling, and mechanical feedback that regulates airway function, it is important to mimic properties of the native ECM such as lung tissue stiffness (0.44-7.5kPa)^48,49^.

#### Optimization of GelMA Hydrogel

In the human conducting airways, epithelial and endothelial cells form monolayers on the ECM whereas fibroblasts are embedded within a 3D interstitial matrix. To replicate this environment, the hydrogel must support both configurations and balance mechanical strength with biological compatibility for AOC integration. It should be injectable and photopolymerizable *in situ*, forming a stable 3D structure that supports pre-vascularization templates and long-term ALI culture. Biologically, it must sustain fibroblast encapsulation while enabling epithelial and endothelial adhesion for co-culture and lumen formation. To identify an optimal GelMA/LAP formulation, nine variants were screened for crosslinking, handling, and fibroblast compatibility. Assessments on viability, morphology, and matrix remodeling were undertaken using GelMA constructs (**Fig. S1)**, to increase throughput and avoid confounding effects from the AOC device materials or continuous flow.

### Crosslinking Efficiency and Density

Across all tested GelMA/LAP formulations, sol fractions were below 40%, indicating successful formation of a hydrogel network upon photopolymerization (**Fig. 1a**). In general, crosslinking efficiency appeared to be dependent on GelMA (**Fig. 1ai**) and LAP photoinitiator (**Fig. 1aii)** concentration; a decrease in sol fraction was observed when GelMA or LAP concentration was increased. When LAP was kept constant, sol fraction gradually decreased from 38.20 ± 0.41 to 10.13 ± 6.64 in 0.6% LAP hydrogels and 32.48 ± 1.60 to 15.58 ± 1.72 in 1.2% LAP hydrogels. When GelMA was kept constant, sol fraction gradually decreased from 26.58 ± 0.62 to 15.58 ± 1.72 in 6% GelMA hydrogels and from 33.33 ± 0.10 to 10.13 ± 6.64 in 12% GelMA hydrogels. No consistent trend was observed in hydrogels composed of 3% GelMA or 0.3% LAP.

**Fig. 1.**
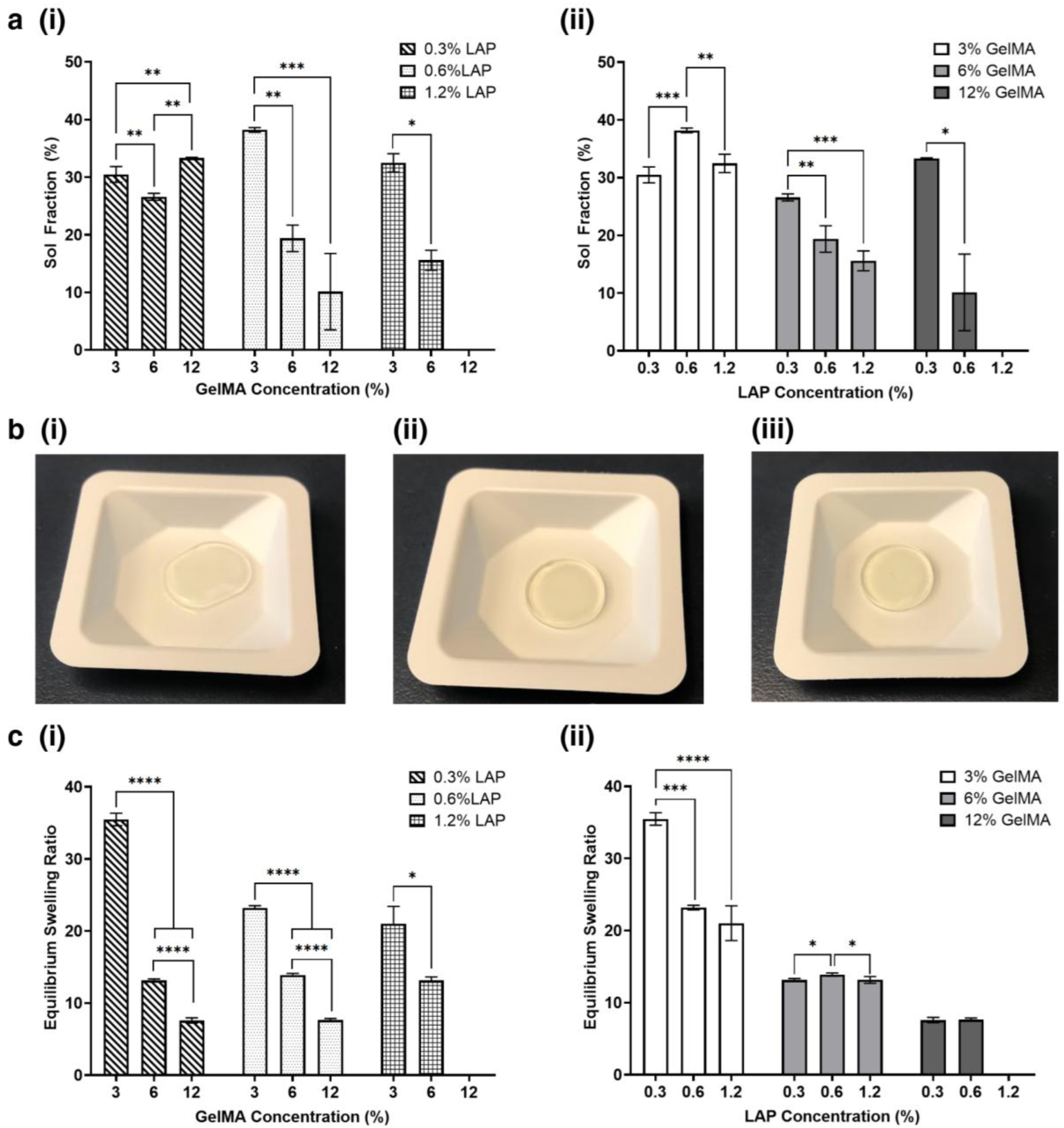
Physical properties of GelMA-based hydrogels with varying GelMA and LAP concentrations. (a) Sol fraction of GelMA formulations. (i) Data plotted by GelMA concentration (3%, 6%, 12%) at different LAP concentrations (0.3%, 0.6%, 1.2%) (ii) Same dataset re-plotted by LAP concentration (0.3%, 0.6%, 1.2%) at different GelMA concentrations (3%, 6%, 12%) to highlight comparative trends. (b) Representative images of hydrogel samples demonstrating that (i) soft 3% hydrogels were able to form a hydrogel but not hold a circular shape when transferred into weighing dish. (ii) 6% and (iii) 12% hydrogels maintained a circular shape after transfer. All hydrogels were made using 0.6% LAP. (c) Swelling ratio of GelMA formulations. (i) Data plotted by GelMA concentration (3%, 6%, 12%) at different LAP concentrations (0.3%, 0.6%, 1.2%) (ii) Same dataset re-plotted by LAP concentration (0.3%, 0.6%,1.2%) at different GelMA concentrations (3%, 6%, 12%) to highlight comparative trends. Data are presented as mean ± standard deviation (n=3). P values calculated using Bonferroni Test. * p < 0.05, ** p<0.01, *** p<0.001, **** p<0.0001.

3% GelMA hydrogels exhibited sol fractions > 30% and were mechanically too soft to maintain their 3D shape (**Fig. 1bi**), causing challenges for both analysis and integration due to sample loss during weighing transfers and poor shape fidelity which compromised the hydrogels’ ability to maintain internal microarchitecture. 6% and 12% hydrogels with 0.3% LAP exhibited sol fractions comparable to 3% GelMA formulations. These hydrogels exhibited reduced shape fidelity in comparison to 6% and 12% hydrogels with higher LAP concentrations (0.6 or 1.2%) which were able to form stable 3D structures (**Fig. 1bii-iii**). Hydrogels composed of 12% GelMA/1.2% LAP were not analyzed as they were highly viscous and therefore difficult to pipette resulting in samples with large bubbles and heterogenous gelation.

Mass swelling ratios for GelMA/LAP hydrogels ranged from 7.58 ± 0.37 to 35.48 ± 0.87 with 3% hydrogels exhibiting the highest swelling ratios and 12% hydrogels exhibiting the lowest (**Fig. 1c**). As shown in **Fig. 1ci**, swelling generally decreased with increasing GelMA concentration indicating higher crosslinking density and reduced porosity when LAP concentration was fixed. Altering LAP concentration had minimal effect on swelling in 6% and 12% GelMA hydrogels indicating constant network density. Combined with sol fraction data, this suggests that increasing LAP can improve shape fidelity without altering the network structure (**Fig. 1cii**). From both a crosslinking and physical perspective, hydrogels composed of 6% GelMA/0.6% LAP, 6% GelMA/1.2% LAP, or 12% GelMA/0.6% LAP demonstrated the most favorable properties and potential for AOC integration.

### Cell Encapsulation

The cell encapsulation capacity of nine GelMA/LAP formulations was assessed using fibroblast-laden hydrogel constructs (**Fig. S1a**). As shown in **Fig. 2ai-iii, b**, fibroblasts encapsulated in 3% GelMA hydrogels were viable (> 80%), but the gel was mechanically too weak to maintain a 3D structure, resulting in 2D cell growth. In contrast, 6% and 12% GelMA hydrogels maintained stable structures allowing cells to grow in 3D, however, viability and morphology of encapsulated cells varied depending on the GelMA and LAP concentration. Fibroblast cell death increased with both higher GelMA (**Fig. 2bi**) and LAP concentrations (**Fig. 2bii**). Denser networks from increased GelMA reduced water sorption capabilities as supported by swelling data (**Fig. 1ci**), thereby reducing cells’ access to nutrients and oxygen^44,50^. Higher LAP elevated levels of free radicals during photopolymerization, which can be cytotoxic^51^. Only 6% GelMA/0.3% LAP and 6% GelMA/0.6% LAP formulations exhibited fibroblast viability above 70%.

**Fig. 2.**
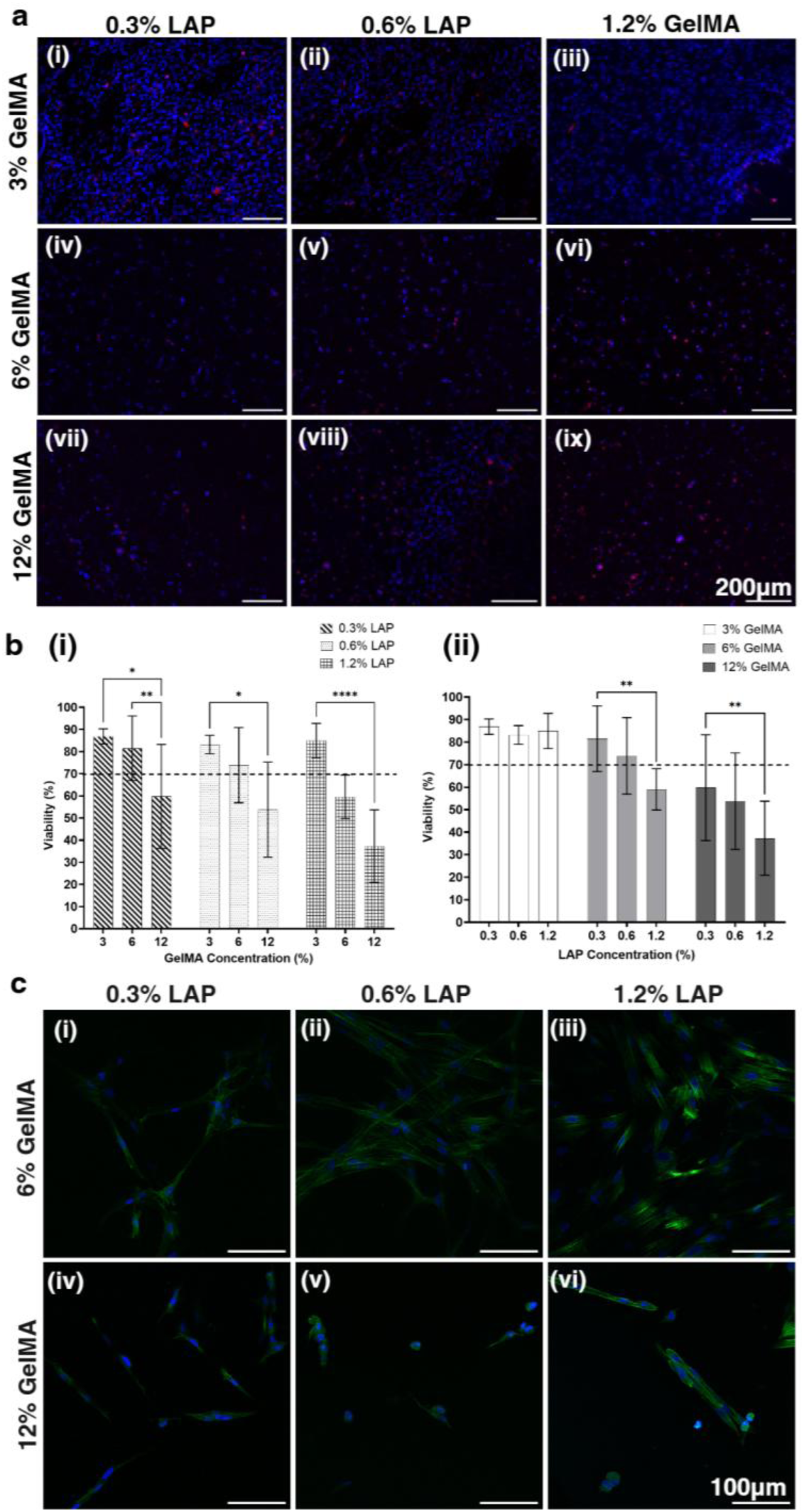
Viability and morphology of encapsulated fibroblasts (a) Representative fluorescence images of fibroblasts encapsulated in GelMA constructs composed of GelMA (3%, 6% and 12%) and LAP (0.3%, 0.6% and 1.2%) after 4 days. Dead cells were stained with Ethidium Homodimer-1 (red) and nuclei were stained with Hoechst 33342 (blue). Each representative image is of a single optical plane. Nuclei staining in (i-iii) 3% GelMA hydrogels was only in a single optical plane, while (iv-vi) 6% GelMA hydrogels and (vii-ix) 12% GelMA hydrogels showed nuclei staining throughout the entire gel. (b) Quantification of viability of encapsulated fibroblasts in GelMA hydrogels composed of different combinations of GelMA (3%, 6%, 12%) and LAP (0.3%, 0.6%, 1.2%). (i) Data plotted by GelMA concentration (3%, 6%, 12%) at different LAP concentrations (0.3%, 0.6%, 1.2%). (ii) Same dataset re-plotted by LAP concentration (0.3%, 0.6%, 1.2%) at different GelMA concentrations (3%, 6%, 12%) to highlight comparative trends. Data presented as mean ± standard deviation. P values calculated using Bonferroni Test. *p < 0.05, ** p<0.01, *** p<0.001, **** p<0.0001. (C) Representative maximum intensity projection (MIP) confocal images of fibroblasts in GelMA hydrogels composed of different combinations of GelMA and LAP. F-actin was stained with Alexa Fluor 488-conjugated phalloidin (green) and nuclei stained with DAPI (blue).

Fibroblasts encapsulated in 6% GelMA hydrogels (**Fig. 2ci-iii)** displayed characteristic elongated morphology and were in closer proximity to one another in comparison to those encapsulated in 12% GelMA hydrogels, facilitating cell-cell communication. This was particularly evident in 6% GelMA/0.6% LAP hydrogels (**Fig. 2cii**) where the fibroblasts appear to form an interconnected network. In contrast, fibroblasts in 12% GelMA hydrogels (**Fig. 2civ-vi)** were more spherical with short extensions and appeared more widely spaced. The 6% GelMA/0.6% LAP formulation provided the best balance between mechanical stability and cell viability, creating a stable 3D structure that supported viable (>70%) cells and enabled them to adopt elongated morphologies, therefore this formulation was selected for further studies and potential integration into the AOC platform.

In addition to cell survival and morphology, the ability to respond to stimuli such as TGF-β1 is essential for capturing stromal cell behaviors and responses such as fibroblast-to-myofibroblast differentiation. Fibroblasts embedded within 6% GelMA/0.6% LAP hydrogel constructs displayed morphological changes when treated with TGF-β1, shifting from an elongated shape to a more star- or web-shaped morphology (**Fig. 3a**). This shift is reflected in the decreased percentage of fibroblasts displaying elongated morphology (**Fig. 3b**). Similarly, the increase in F-actin staining (**Fig. 3c**) in TGF-β1 treated cells reflects the increase in cytoplasmic actin stress fibers present in myofibroblasts. The morphological shift coupled with increased alpha smooth muscle actin (α-SMA) expression (**Fig. 3d**) indicates the presence of myofibroblasts. Together, these results confirm that fibroblasts in the 6% GelMA/0.6% LAP hydrogel successfully undergo fibroblast-to-myofibroblast differentiation indicating that this formulation does not hinder fibroblast function nor differentiation capacity and supports its integration into the AOC platform.

**Fig. 3.**
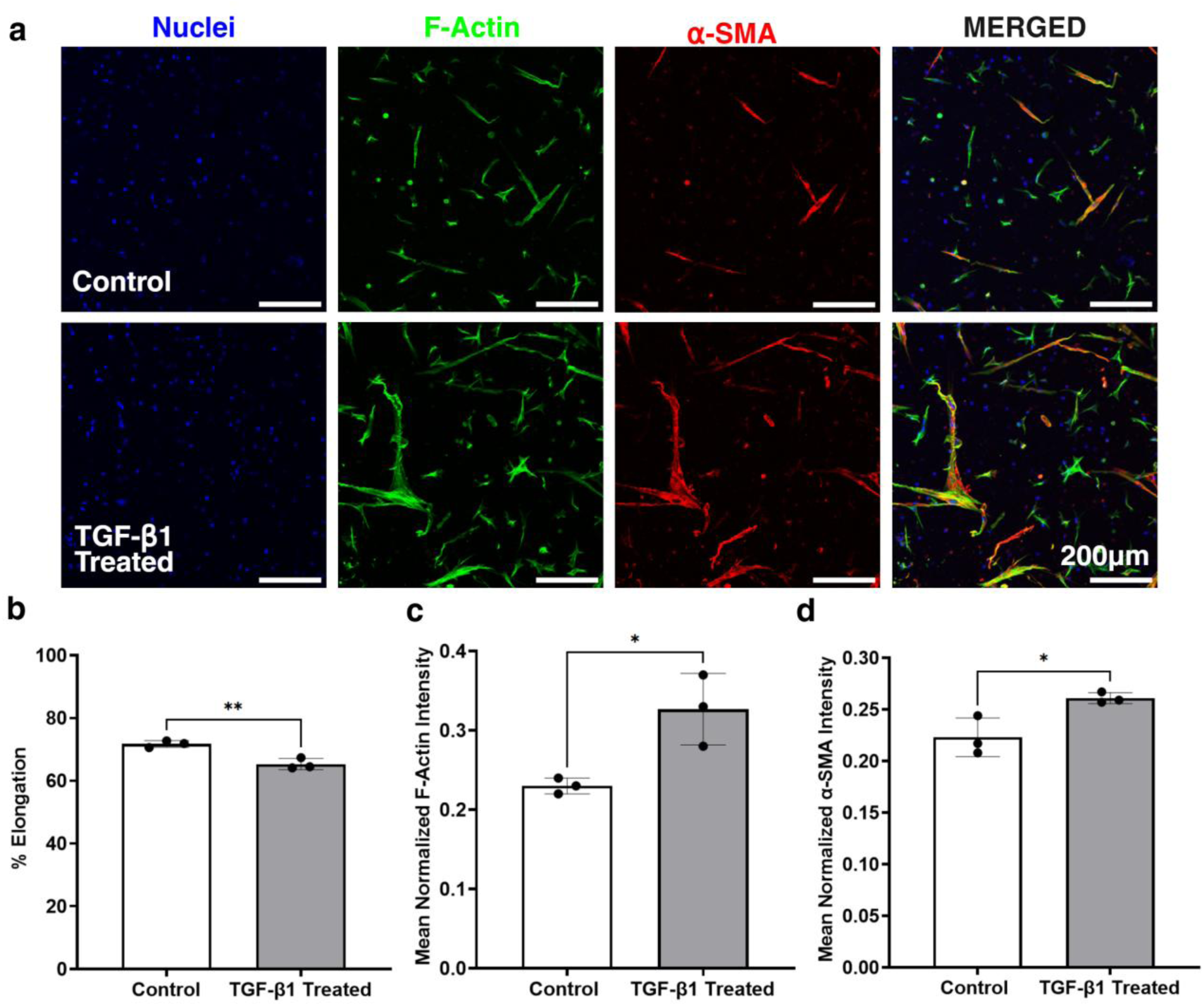
TGF-β1 treatment in 6% GelMA/0.6% LAP hydrogel constructs. Lung fibroblasts (HFL1) were encapsulated within hydrogels composed of 6% GelMA/0.6% LAP treated with DMEM/F-12 cell culture medium plus 10% FBS without (control) and with TGF-β1 (50 ng/mL) for 72 hours. (a) Representative MIP confocal images of HFL1 cells encapsulated in 6% GelMA/0.6% LAP constructs. F-actin (green), α-SMA (red) and nuclei (blue) were stained to visualize cell morphology, fibroblast activation, and myofibroblast differentiation. Images were captured at 10x. (b) Quantification of percentage of fibroblasts displaying elongated morphology (elongation). Elongation decreased with TGF-β1 treatment indicating a transition away from fibroblast morphology. (c) Quantification of mean intensity of F-actin in samples normalized to total cells. The statistically significant increase seen with TGF-β1 treatment indicates an increase in cytoplasmic actin stress fibers. (d) Quantification of mean fluorescence intensity of α-SMA positive cells normalized to total cells. The statistically significant increase seen with TGF-β1 treatment indicates fibroblast activation. Data represents mean ± standard deviation (n=3). P values calculated using two tailed t-test. * p < 0.05 and **p<0.01.

### Cell Adhesion

Beyond fibroblast encapsulation, cell attachment, spreading, and growth on the hydrogel surface are essential for establishing tissue interfaces within the AOC, particularly at the stromal-endothelial boundary. Both endothelial and epithelial cells were able to adhere, proliferate, and form confluent monolayers on the surface of 6% GelMA/0.6% LAP hydrogels, demonstrating its positive cell-binding behavior and suitability for integration. Human umbilical vein endothelial cell (HUVECs) (**Fig. 4a)** displayed characteristic cobblestone morphology and a monolayer which for AOC integration is essential as the lumen-patterned hydrogel is used as the endothelial substrate and monolayer formation is the basis for recreating the stromal-vascular interface. Epithelial cells (**Fig. 4b)** also showed cobblestone morphology and a thick cell monolayer. For both cell types, the hydrogel’s surface functions as a basement membrane surrogate. The GelMA hydrogel’s intrinsic cell adhesivity supports surface-bound cell layers without additional ECM components, and co-culture capacity (**Figure. S2**), providing an environment capable of capturing physiologically relevant cell-cell and cell-ECM interactions, making it suitable for integration into the AOC.

**Fig. 4.**
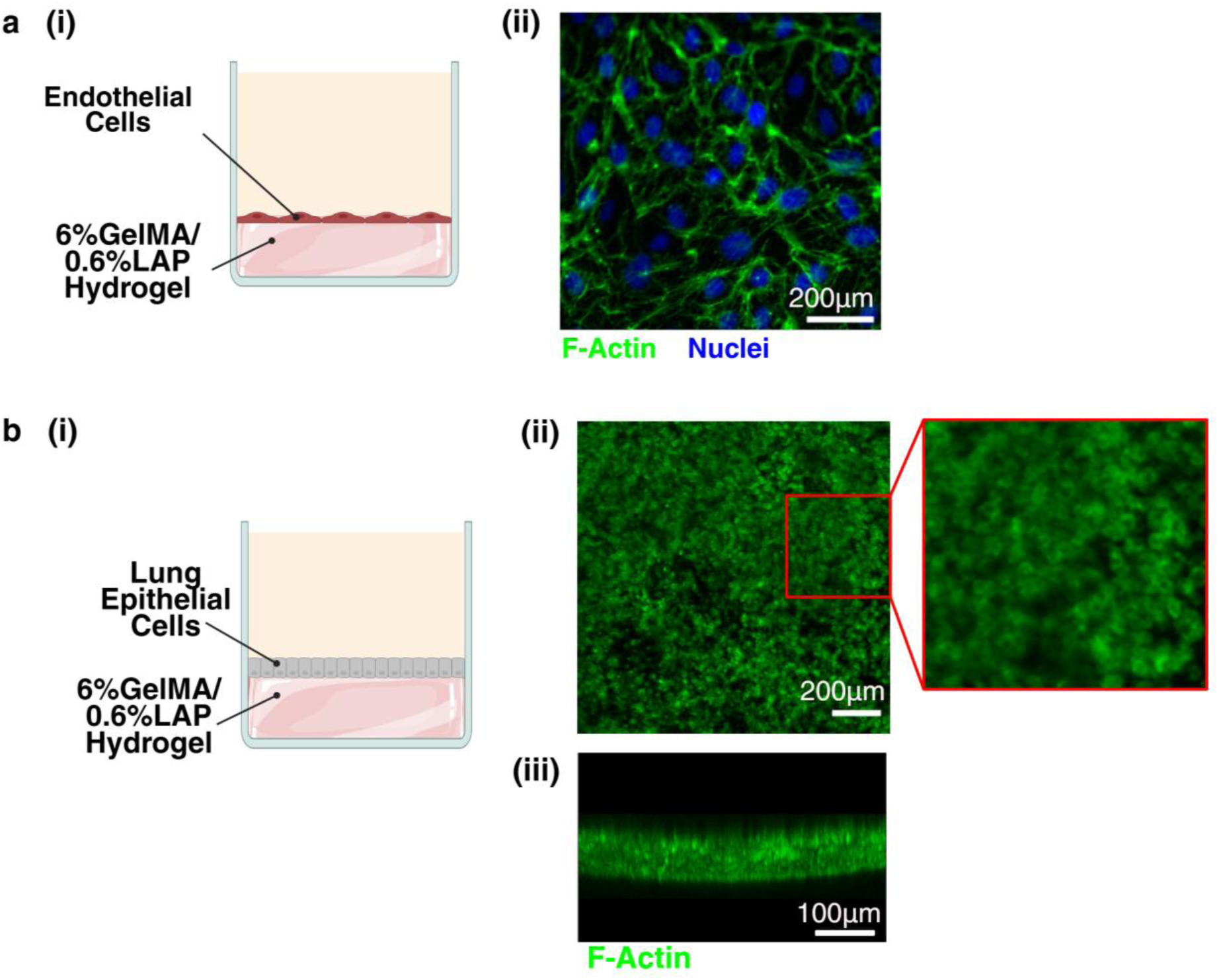
Cell adhesion on 6% GelMA/0.6% LAP hydrogel surface (a) Human umbilical vein endothelial cells (HUVECs) cultured on GelMA hydrogel surface. (i) Schematic of HUVECs seeded on GelMA hydrogel construct. (ii) Representative MIP confocal image highlighting cobblestone morphology. HUVECs were stained with Alexa Fluor 488-conjugated phalloidin (green) and Hoechst 33342 (blue). Merged image of fluorescent channels captured at 40x. (b) Airway epithelial cells (Calu-3) cells cultured on GelMA hydrogel surface. (i) Schematic of Calu-3 cells seeded GelMA hydrogel construct. (Bii-iii) Representative MIP confocal image showing epithelial monolayer formation. Calu-3 cells were stained with Alexa Fluor 488-conjugated phalloidin and images captured at 10x. (ii) The representative MIP of the confluent monolayer with inset highlighting cobblestone morphology. (biii) The 3D reconstruction (cross-section) highlights the cell monolayer thickness. Schematics created with BioRender.com.

### Developing Airway-On-A-Chip with Integrated GelMA hydrogel

#### Device Design and Fabrication

The AOC (**Fig. 5**) combines a PDMS-based microfluidic platform with a patterned GelMA hydrogel to more accurately recapitulate the 3D structural and functional organization of the human small conducting airways. The device comprises two chambers with three spatially distinct and independenly accessible cell compartments; an upper microchannel, 3D hydrogel, and lumen. As illustrated in **Fig. 5a-b**, two microchannels containing PDMS layers and a thin porous membrane combine to form the external “housing” device that accomodates the hydrogel component and facilitates lumen patterning after assembly. When assembled, the microchannels vertically stack, creating a central fluidic channel. The membrane sandwiched between the two PDMS layers operates as a culture surface (≈ 18 mm^2^) and a semi-permeable partition. PDMS was selected for its ease of microfabrication, biocompatibility, and gas permeability. Guided by anatomical dimensions and separations, the top chamber was designed to represent the luminal “air-facing side” of the conducting airway, while the bottom chamber was designed to represent the basal “tissue-facing” side containing an interstitial ECM hydrogel and perfusable vasculature-like lumen. The top microchannel has a hydraulic diameter of 1 mm reflecting the diameter of a human bronchiole, similar to other airway LOC models^24,26–28,30,31,33,37,38^.

**Fig. 5.**
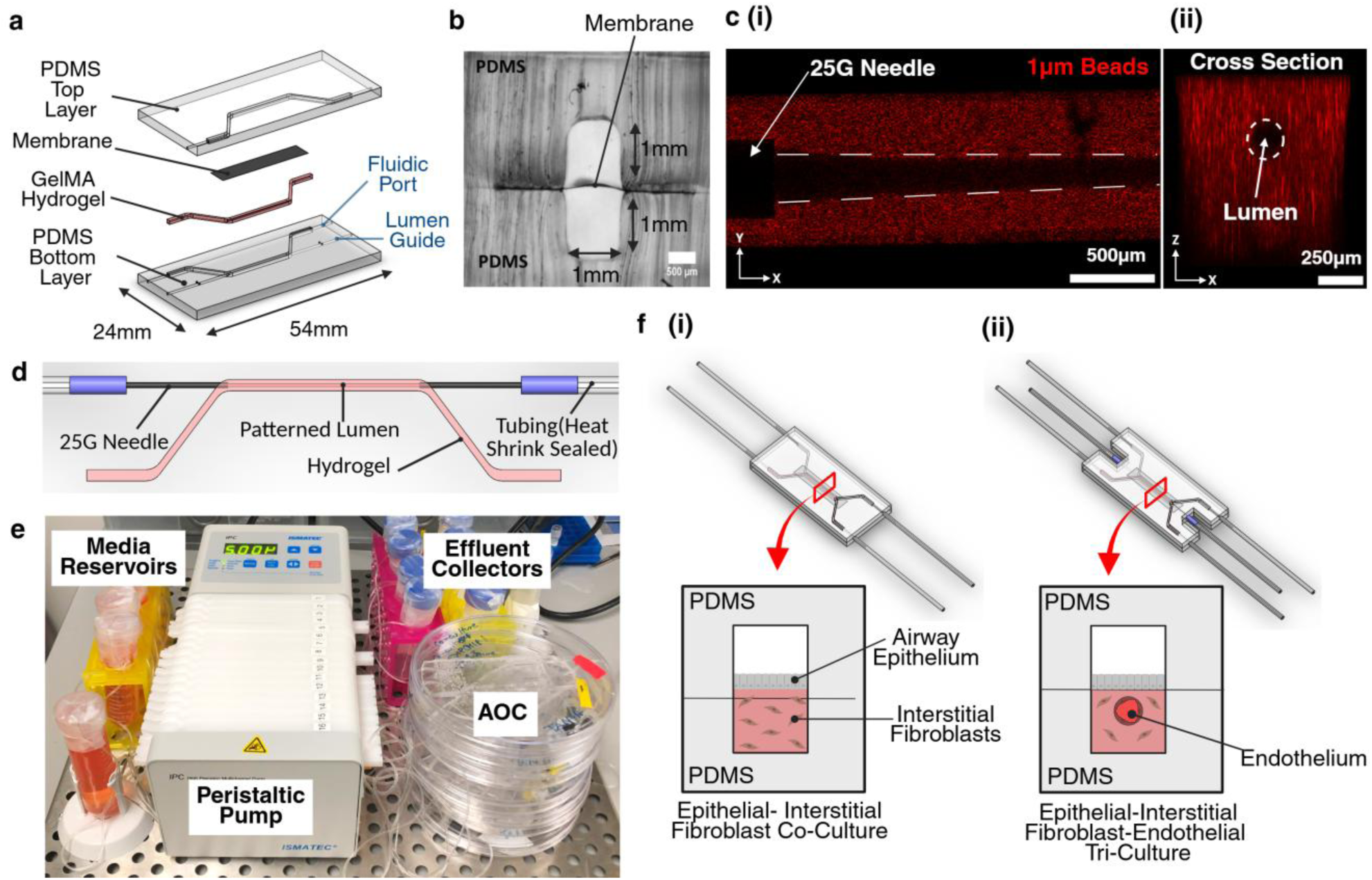
Airway-on-a-chip microfluidic device. (a) Exploded view of the layers that make up the microfluidic device and integrated hydrogel. (b) Brightfield microscopy image of a cross section of AOC microdevice showing PDMS enclosed channels separated by a porous membrane creating two independent compartments. (c) Lumen structure patterned within 6%GelMA/0.6% LAP hydrogel in multilayered AOC microdevice. Red (542 nm) fluorescent microbeads show the hollow 3D microarchitecture that results from patterning. The hydrogel pre-polymer solution loaded and polymerized in the bottom microchannel is mixed with microbeads for visualization. (i) Representative MIP illustrating how the lumen sits within the bottom microchannel and connects to a needle for fluid introduction. (ii) 3D confocal reconstruction image illustrating circular lumen geometry with hydrogel surrounding the lumen structure. (d) Illustration of fluidics connections to lumen directly within the hydrogel contained within the bottom microchannel. (e) AOC microdevices connected to a peristaltic pump-based perfusion system. Connection to this system provides a continuous supply of cell culture media and mimics hemodynamic shear stresses. (f) Schematic of assembled AOC microdevice showing possible cell culture layouts. (i) Epithelial-interstitial fibroblast co-culture (ii) Epithelial-interstitial fibroblast-endothelial tri-culture. Removing media from the upper microchannel and coupling with the airflow system enables epithelial cells to be exposed to air that can trigger differentiation of airway epithelial cells and mimic luminal shear stresses. Created with BioRender.com

The bottom microchannel confines the GelMA pre-polymer, enabling controlled *in situ* photopolymerization and lumen patterning. The optical transparency of PDMS and separation membrane facilitates precise gel loading and *in situ* photopolymerization. The microchannel walls and separation membrane prevent leakage of the liquid pre-polymer into the upper microchannel preserving compartment integrity. Embedded lumen guides enable compatibility with sacrificial molding techniques and repeatable lumen positioning with minimal device modifications. Needle positioning was optimized according to physiological oxygen diffusion limits (100-200µm^52^), enabling the device to be operated at an air-tissue interface without comprising the ability to supply cells with sufficient oxygen and nutrients via lumen perfusion. Cells can be easily incorporated into the GelMA pre-polymer, loaded around the removable acupuncture needle and rapidly polymerized *in situ* upon irradiation with violet light (399 nm), forming a stable 3D scaffold with cells distributed throughout. Using a photopolymerizable GelMA hydrogel enables ease of gel loading and *in situ* polymerization, both of which facilitate cell encapsulation, delivery of the hydrogel to a specific location with high spatial resolution, and lumen patterning. The needle used as the sacrificial template defines the lumen size and geometry, producing a circular vessel-like lumen (∼250 µm diameter) within the hydrogel that can be immediately perfused after photopolymerization and needle removal to support the cell culture. The strong covalently bonded network that forms upon photopolymerization^44^ exhibits high shape fidelity and mechanical stability.

Simultaneous formation of the 3D hydrogel and lumen creates two distinct yet interactive microenvironments within the subepithelial compartment of the AOC, enabling direct cell-cell communication and cell-ECM interactions. **Fig. 5c** illustrates the subepithelial connective tissue-like structures established within the AOC, highlighting the circular vasculature-like lumen spanning the length of the 3D interstitial ECM analog contained within the central flow region of the bottom microchannel (**Fig. 5ci**). The lumen operates as a secondary fluidic channel within the AOC as the hydrogel circumferentially surrounds it (**Fig. 5cii**) and the lumen openings interface directly with the fluidic connections (**Fig. 5d**). The circular profile promotes uniform shear, an advantage not achievable in models using rectangular microchannels^53^.

Integration with perfusion (**Fig. 5e**) and airflow systems (**Fig. S3.2**) enables precise control over the dynamic microenvironments estbalished within each compartment. Continuous perfusion supplies oxygen and nutrients while removing waste, while the custom airflow system delivers filtered air tangentially across the apical surface of the epithelial layer introducing realistic airflow-induced shear stresses known to enhance epithelial function and differentiation^28,32,33^. Unidirectional airflow was used to model inhalation in proof-of-concept exposure studies. Positioning the vacuum-driven pump downstream of the AOC minimizes contamination risk and enables coupling with the whole wood smoke exposure system (described in **S3** and illustrated in **Fig. S3.1, S3.2**). This provides a controlled way to introduce smoke and conduct realistic aerosol exposures while preserving the established cell culture and ALI conditions. The porosity and gas permeability of GelMA^54–56^ further supports diffusion of smoke-derived gases into the various tissue compartments. As illustrated in **Fig. 5f** the AOC can be operated in multiple configurations such as an epithelial-fibroblast co-culture with unpatterned cell-laden hydrogel (**Fig. 5fi**) or tri-culture incorporating epithelial, fibroblast and endothelial cells (**Fig. 5fii**). Both configurations support compartmentalized yet interconnected cellular microenvironments.

#### Establishing An Epithelial-Fibroblast-Endothelial Tri-Culture Within Airway-On-A-Chip

In the native bronchioles, epithelial, stromal, and endothelial cells exist in a highly organized 3D microenvironment and interact dynamically through the ECM. The AOC replicates this organization by seeding the upper “epithelial” and lower subepithelial “stromal” compartment to create an epithelial-fibroblast-endothelial tri-culture (**Fig. 6a-b**)

**Fig. 6.**
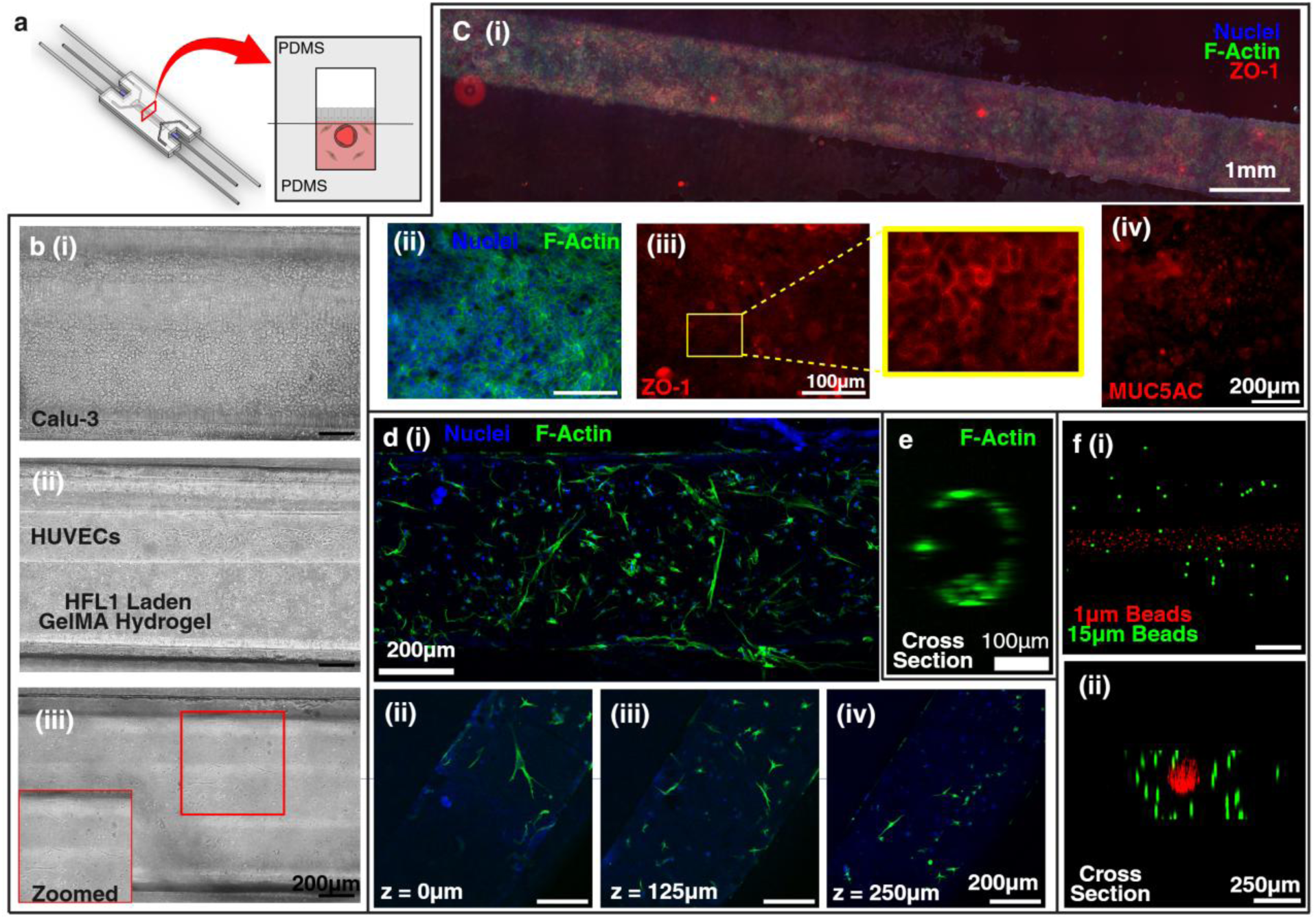
Airway-on-a-chip containing lumen patterned-fibroblast-laden GelMA hydrogel (a) Schematic of AOC in epithelial-interstitial fibroblast-endothelial tri-culture format highlighting the three independent cellular compartments that can be seeded with airway epithelial cells, lung fibroblasts and microvascular endothelial cells. Created with BioRender.com. (b) Brightfield images showing (i) Calu-3 cells on the membrane surface within the top microchannel and (ii-iii) the endothelial-seeded lumen patterned in the fibroblast-laden 6%GeMA,0.6%LAP hydrogel contained within the bottom microchannel. (ii) Image captured at the center of the lumen structure highlighting how HUVECs are confined to the lumen, while HFL1 cells are distributed throughout the surrounding hydrogel. (iii) At the interface of the top and bottom microchannels, the lumen cannot be seen highlighting that the fibroblast laden hydrogel envelops the lumen structure. The inset highlights the elongated morphology of encapsulated HFL1 cells. (c) Calu-3 cells grown within AOC. (i) Stitched image of extracted membrane displaying an epithelial monolayer that runs the entirety of the microchannel. Immunofluorescence images captured at 10x of Calu-3 cells stained with (ii) DAPI (blue) and Alexa Fluor 488-conjugated Phalloidin (green) to visualize F-actin and nuclei and (iii) ZO-1 (red) to visualize tight junctions and illustrate cobblestone morphology. In the inset, clear, smooth borders of ZO-1 can be seen. (iv) Immunofluorescence images captured at 20x of Calu-3 cells highlighting the presence of mucin-producing goblet cells (MUC5AC). MUC5AC staining appears as small dots. (d) HFL1 cells encapsulated in GelMA hydrogel extracted from AOC microdevice. (d) F-actin (green) and nuclei (blue) staining visualizes fibroblast cell distribution within the 6% GelMA/0.6%LAP hydrogel. (dii-iv) Confocal slices at various depths demonstrate elongated morphology throughout the hydrogel, confirming its 3D structure. (e) Endothelial lined lumen in GelMA hydrogel extracted from AOC. (f) Spatial arrangement of lumen within fluorescent bead laden hydrogel from a (i) top view and (ii) side view demonstrates that components embedded within the GelMA bulk circumferentially surround the lumen structure.

### Epithelial Compartment

Airway epithelial cells (Calu-3) seeded in the top microchannel adhered to the surface of the ECM protein-coated porous membrane that served to mimic the basement membrane, forming a continuous epithelial monolayer spanning the entire length of the microchannel (**Fig. 6ci**) that exhibited key features of the airway epithelium. Exposure to air via the integrated airflow system enabled the AOC to mimic the air-tissue interface and function of the conducting airways. The epithelial monolayer displays characteristic cobblestone morphology (**Fig. 6cii**) and ZO-1 positive and MUC5AC positive staining, which indicates the presence of tight junctions and goblet cells, respectively (**Fig. 6ciii-iv**). Together, these findings demonstrate that the AOC supports formation of an epithelium that mimics airway barrier function and integrity, while also enabling differentiation towards a mucus-producing phenotype. Although Calu-3 cells do not fully recapitulate the diversity of primary airway epithelium—lacking ciliated and club cells—the presence of a robust, mucus-producing epithelial barrier within the AOC demonstrates, via proof-of-concept, the potential of the AOC for modeling the structure and function of the epithelium that lines the airway.

### Subepithelial Compartment

The lung fibroblast-laden 6% GelMA/0.6% LAP hydrogel provided a physiologically relevant 3D environment mimicking the interstitial ECM and resident fibroblasts (**Fig. 6d**). Fibroblasts maintained elongated morphologies consistent with healthy stromal cells (**Fig. 6biii**). Needle-based patterning produced a perfusable lumen within the hydrogel without compromising encapsulated fibroblasts’ ability to grow. Using fluorescent beads to represent the fibroblasts and endothelial cells (**Fig. 6f**), demonstrated the stromal-vascular interface created in the subepithelial compartment which enables enhanced cell-cell crosstalk and cell-ECM interactions.

Endothelial cells (HUVECs) seeded into the lumen post-patterning utilized the lumen’s internal surface as a vessel template, attaching to the hydrogel in a 3D manner and forming circular vascular-like geometry (**Fig. 6e**) which is important for recapitulating the endothelium^57–61^. While HUVECs originate from large vessels and lack tissue specific microvascular characteristics, their successful integration into AOC validates the platform’s vascularization capacity and establishes a proof-of-concept for tri-culture modeling.

### Proof-Of-Concept Studies to Demonstrate AOC Utility for Exposure Studies

#### On Chip TGF-β1 Treatment to Elicit Inhalation Response

In addition to reproducing healthy airway architecture, a physiologically relevant model must capture biological responses associated with airway remodeling and smoke exposure. To demonstrate the AOC’s utility, fibroblast-laden devices were treated with TGF-β1 and whole wood smoke. TGF-β1, a central regulator of lung function and fibrosis, drives fibroblast-to-myofibroblast differentiation, ECM deposition, and inflammatory signaling.

TGF-β1 treatment on a monoculture AOC (**Fig. 7b-e**) induced clear morphological changes in fibroblasts. Untreated fibroblasts maintained an elongated morphology, whereas treated cells adopted a star- or web-like shape, consistent with activation. This transition was accompanied by an increase in fibroblast number and early evidence of myofibroblast differentiation, including α-SMA expression and stress fiber formation (**Fig. 7b-d**). Although the response was less pronounced than in static GelMA cultures (**Fig. 3, S2**), the AOC successfully recapitulated TGF-β1-induced activation and differentiation. The reduced response likely reflects reduced cytokine contact time and cytokine dilution due to perfusion and adsorption in the tubing. Adjusting cytokine concentration or flow conditions may enhance differentiation while maintaining the platform’s physiological relevance.

**Fig. 7.**
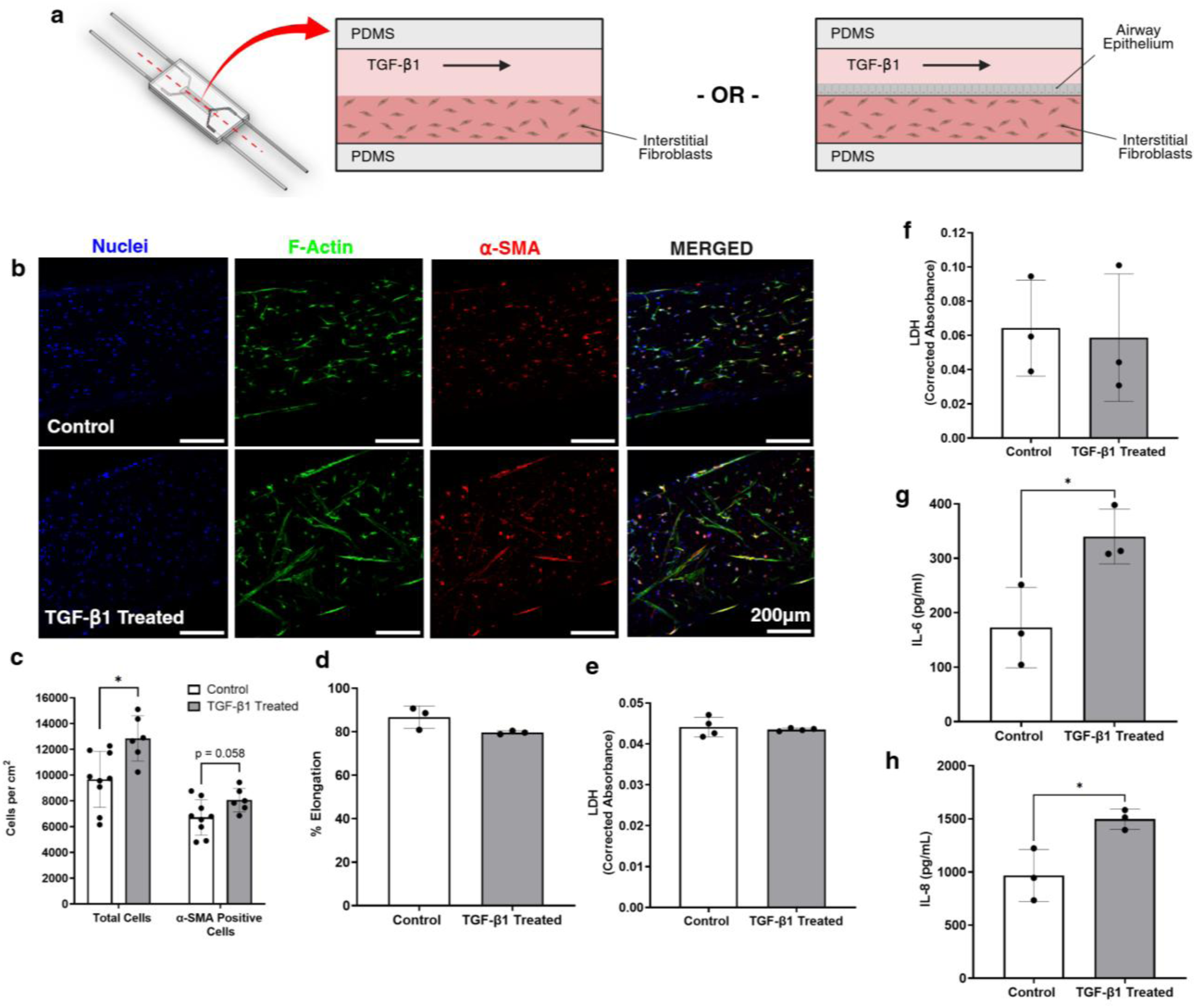
On-chip TGF-β1 treatment in AOC (a) Schematic of AOC interstitial fibroblast monoculture and epithelial-interstitial fibroblast co-culture configurations used for treatments. During treatment, media containing TGF-β1 (50 ng/ml) is continuously perfused through the top microchannel. Control AOCs were perfused with a fresh volume of serum-reduced culture media (1% FBS) under the same conditions for comparisons. Created with BioRender.com. (b-e) TGF-β1 treatment in interstitial fibroblast monoculture AOC to induce fibroblast activation and myofibroblast differentiation. (b) Representative MIP confocal images of fibroblasts encapsulated in 6%GelMA/0.6%LAP hydrogel containing AOC. Cells were stained with Hoechst (Nuclei; blue), Alexa Fluor 488-conjugated phalloidin (F-actin; green) and α-SMA (red). (c) Quantification of total number of fibroblasts (total cells) within the AOC model in comparison to the number of cells expressing α-SMA. Data represents mean ± standard deviation (n=3). For each sample, a z-stack (25 µm step size) at three regions of interest for controls and two regions of interest for treated chips were acquired using a 10x objective. * p < 0.05. (c) Quantification of fibroblast elongation. Data represents mean ± standard deviation (n=3). (e) Quantification of LDH release in effluent collected from the AOC model. Data represents mean ± standard deviation (n=4). (f-h) Secretion from TGF-β1 treated epithelial-interstitial fibroblast co-culture AOCs (f) LDH levels show no significant indicators of cytotoxicity (g-h) Cytokine levels in effluent collected from AOC models. Increased secretion of (g) IL-6 and (h) IL-8 demonstrates that the co-culture AOC model can mimic an inflammatory response to TGF-β1. Data are presented as mean ± standard deviation (n=3). P values calculated using two tailed t-test. * p < 0.05.

In co-culture AOCs (**Fig. 7f-h**), TGF-β1 treatment induced a significant increase in interleukin 6 (IL-6) and 8 (IL-8) secretion without evidence of cytotoxicity. These cytokines are central mediators of airway inflammation and remodeling. Upon TGF-β1 stimulation, fibroblasts and myofibroblasts are known to release IL-6 and IL-8, while airway epithelial cells release IL-6 but may inhibit IL-8 secretion^62,63^. These results demonstrate that the AOC can recapitulate cytokine responses associated with elevated TGF-β1, supporting its utility for modeling early fibrotic and inflammatory processes.

#### On Chip Exposure to Whole Wood Smoke

The airways serve as the primary interface between the external environment and the body, making them vulnerable to inhaled particulates and pathogens that can disrupt cellular function, interactions, and responses. Previous *in vitro* studies using wood smoke extracts have shown that wood smoke can be cytotoxic to cells^64,65^, and exposure can induce oxidative stress, epithelial barrier dysfunction, ECM remodeling and inflammation^65–67^. Developing physiologically relevant *in vitro* models that can replicate these exposures is critical for studying inhalation-induced injury and inflammation, particularly as wildfire smoke poses increasing health risks. To address this, the AOC was integrated with a smoke exposure system containing a smoke generator (**Fig. S3.1, S3.2**) to simulate wildfire smoke exposures. The system enabled direct exposure to whole wood smoke with controlled PM2.5 delivery at an air-tissue interface under realistic airflow. The AOC’s vertical stacked microfluidic configuration orients cell layers perpendicular to airflow (**Fig. 8a**) allowing smoke to flow tangentially across the epithelium, mimicking luminal shear stress and gravity-driven particulate deposition^30^.

**Fig. 8.**
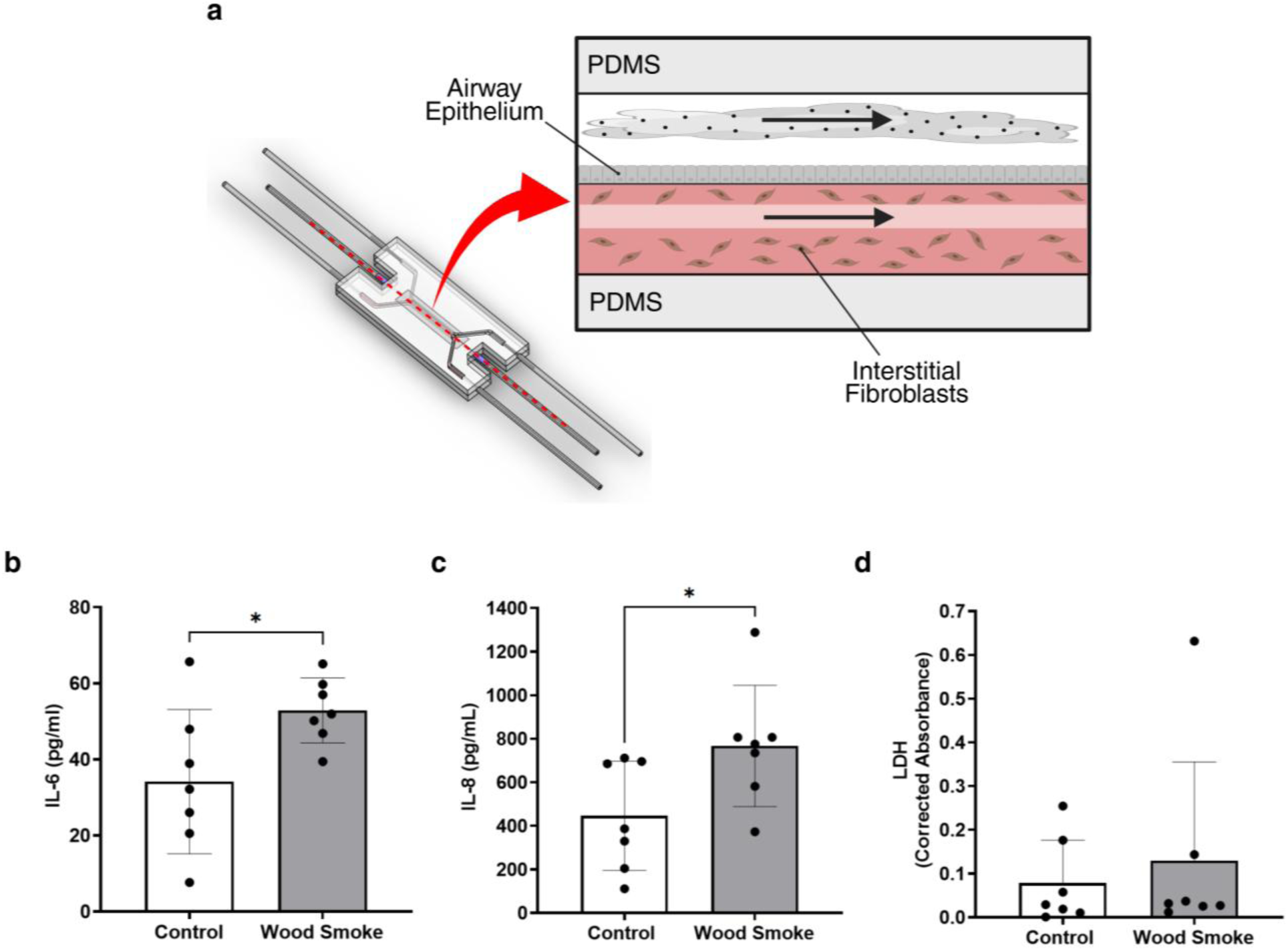
On-chip wood smoke exposure in AOC in epithelial-lumen-patterned interstitial fibroblast co-culture format. (a) Schematic of wood smoke exposure conducted in an AOC. During exposure, aerosolized wood smoke flows through the top microchannel containing an airway epithelium which lies above interstitial fibroblasts embedded throughout an ECM hydrogel. Created with BioRender.com. (b-d) Cytotoxicity and inflammatory mediator release associated with on-chip wood smoke exposure. (b-c) Quantification of cytokine release in effluent collected from AOC containing an epithelial-interstitial fibroblast co-culture exposed to filtered air (control) and wood smoke. AOC models exposed to wood smoke exhibit statistically significant increases in (b) IL-6 and (c) IL-8. (iii) LDH in effluent does not show a significant increase. Data represents mean ± standard deviation (n=7). P values calculated using two tailed t-test. * p < 0.05.

As shown in **Fig. 8b-c**, exposure to wood smoke led to elevated levels of IL-6 and IL-8 in AOC effluent compared to filtered-air controls, consistent with findings from previous human and animal studies^68–73^. Although no cytotoxic effects were observed at the exposure levels used for proof-of-concept studies (**Fig. 8d**), the smoke exposure system allows for controlled increases in PM2.5 concentration and exposure duration to model the cellular damage associated with fine particulate inhalation. Higher PM2.5 doses can be achieved by adjusting the amount of biofuel burned per exposure, with real-time monitoring ensuring precise dose control.

## Discussion

Although hydrogels in general offer several favorable properties for cell culture^50,74–76^, for effective integration into the AOC, the hydrogel must balance mechanical stability, permeability and biological functionality to recapitulate the native airway ECM while satisfying the engineering demands of the microfluidic platform. While natural hydrogels are highly biocompatible but mechanically weak and synthetic hydrogels are robust but less bioactive, GelMA provides a tunable alternative, allowing optimization as demonstrated when used to mimic the alveolar basement membrane from a microstructure and stiffness perspective^17^. Of the formulations tested, 6% GelMA/0.6% LAP best satisfied the AOC requirements, exhibiting effective crosslinking, shape fidelity compatible with lumen patterning, nutrient and oxygen diffusion, and no adverse effects on cell survival, activity or cell-ECM interactions.

Although previous tri-culture airway LOC models^23,25,31,35–37,39^ have contributed valuable insights, they often lack important airway features such as 3D ECM, perfusable circular vasculature, physiological flows, diffusion-based nutrient and oxygen transport and gravity-driven particulate deposition. The absence of these features limits their suitability for airway modelling, particularly in relation to exposure studies. Here, the developed AOC addresses these limitations by integrating a lumen-patterned, cell-laden hydrogel within a membrane-separated microfluidic device, enhancing physiological relevance, and enabling multicellular organization and complex cell-cell and cell-ECM interactions. This configuration supports an air-tissue interface with airflow to mimic luminal shear, and allows 3D spatial arrangement of epithelial, fibroblast and endothelial layers, closely mirroring airway tissue architecture where the epithelium resides above a fibroblast-rich stroma supported by vasculature.

Unlike previous membrane-separated^24,33,37,38^, or 2D-hydrogel-based designs^26,28,32^ the photopolymerizable GelMA hydrogel recreates the small conducting airway’s interstitial ECM and resident fibroblasts, enabling dynamic cell-ECM interactions. The hydrogel’s biodegradability due to MMP-sensitive sites allows encapsulated fibroblasts to remodel its structure^77^, further enhancing the AOC’s ability to simulate airway remodeling and fibrosis.

Sacrificial molding creates a hollow, perfusable vessel-like lumen within the hydrogel, that mimics the vasculature and stromal-vascular interface when seeded with endothelial cells, while allowing realistic nutrient flow and blood flow-related shear stress. The mechanically stable hydrogel supports this internal architecture, re-establishing a dual-channel system where airflow and vascular perfusion are simultaneously simulated. Although current lumen sizes are constrained by template diameter, smaller sacrificial templates (e.g., glass capillaries) or self-vascularization strategies^78^ could yield microvasculature-sized lumens. Embedding endothelial cells alongside fibroblasts during hydrogel photopolymerization could promote spontaneous self-organized vessel formation in response to biological, or mechanical stimuli^78^. Existing pulmonary vasculature-on-a-chip models have examined endothelial mechanotransduction^59,60,79^, matrix remodeling^79^ and pulmonary vasculature diseases^80–83^, but few incorporate airway epithelium or airflow, limiting their suitability for exposure studies. Varone et al.^23^ advanced alveolar modeling through integrating a collagen hydrogel with a membrane-separated microfluidic design. However, as their focus was on the epithelial-stromal interface, their vasculature remained confined to a non-physiological rectangular microchannel and separated from the hydrogel environment by a porous membrane, thereby restricting realistic endothelial-stromal interactions.

On-chip TGF-β1 treatment validated the AOC’s responsiveness, highlighting its utility for recapitulating tissue remodeling, fibrosis and inflammation associated with exposure to particulate matter and smoke. Controlled delivery of biochemical cues through perfusion allows investigation of cytokine-driven fibrotic and inflammatory signaling that is relevant to smoke-induced injury.

Combining the AOC with a whole smoke exposure system enables physiologically relevant aerosol exposure, offering a more realistic alternative to static ALI or extract-based exposures. Real-time smoke generation and monitoring allows precise control over exposure parameters, enabling dynamic studies of cell-particle interactions. The system can be expanded to simulate diverse wildfire smoke profiles by varying biofuel type, PM2.5 dose, burning conditions, and exposure duration. This flexibility would allow investigation of how geographical variations in wildfire emissions can elicit different biological responses. Additionally, controlled dilution offered by the exposure system enables dose-response and temporal smoke profile exposure studies. Whole wood smoke exposure demonstrated the AOC’s utility to replicate a smoke-induced pro-inflammatory response. Further AOC studies should assess smoke-induced barrier dysfunction, myofibroblast differentiation, cytokine profiles (e.g. TGF-β1, TNF-α), and ECM protein deposition to confirm the platform’s robustness.

## Conclusion

In this study, we developed a physiologically relevant AOC integrating a tunable GelMA hydrogel. The microfluidic design, combined with the optimized 6% GelMA/0.6% LAP hydrogel provided the mechanical stability, biocompatibility, and permeability necessary for the AOC to recapitulate structural and functional features of the small conducting airways, including an epithelium, fibroblast-rich 3D interstitial ECM and perfusable vasculature, within a platform capable of controlled dynamic aerosol exposure to wood smoke. Proof-of-concept studies demonstrate that this AOC recapitulates key biological responses associated with airway remodeling and smoke exposure. Together, these findings establish the AOC as an advancement in physiologically relevant *in vitro* airway models for realistic inhalation-based aerosol exposure studies.

Beyond its current application, the photopolymerizable, tunable, and biodegradable nature of GelMA provides the foundation for future AOC advancements. Future AOC platforms can further exploit GelMA to mimic fibrotic or disease-specific microenvironments, as well as eliminate the reliance on the separation membrane. By tuning GelMA stiffness, the AOC could reflect the fibrotic ECM (∼17 kPa)^84,85^ associated with chronic exposure to air pollutants and chronic airway diseases such as COPD^86^, enhancing its utility for exposure studies and disease modeling. Eliminating the synthetic separation membrane would allow the epithelial cell layer to be anchored directly to the underlying lumen patterned, fibroblast-laden hydrogel, enabling more realistic basement membrane modeling. This enhancement could be supported through the addition of micro-anchors for hydrogel stabilization^28^ and a biodegradable membrane for temporary channel separation. A fully hydrogel-based interface would more accurately replicate the epithelial-stromal and stromal-vascular interfaces, expanding the model’s utility for investigating smoke-induced effects on barrier integrity, tissue remodeling, inflammation, and cell-cell and cell-ECM interactions such as epithelial-to-mesenchymal transition^87,88^.

## Methods

### Hydrogel Fabrication

GelMA (50% Degree of Functionalization) and lithium phenyl-2,4,6-trimethylbenzoylphosphinate (LAP) were purchased as lyophilizate and powder, respectively, from CELLINK (BICO). GelMA/LAP solutions were prepared, following the manufacturer’s recommendations. Three GelMA concentrations (3%, 6% and 12% w/v) combined with three LAP concentrations (0.3%, 0.6% and 1.2% w/v) to prepare nine different GelMA/LAP formulations; 3% GelMA/0.3% LAP, 3% GelMA/0.6% LAP, 3% GelMA/1.2% LAP, 6% GelMA/ 0.3% LAP, 6% GelMA/0.6% LAP, 6% GelMA/1.2% LAP, 12% GelMA/ 0.3% LAP, 12% GelMA/ 0.6% LAP and 12% GelMA/1.2% LAP.

Sterile GelMA and LAP were each dissolved separately in Dulbecco’s Modified Eagle’s Medium/Nutrient Mixture F-12 (DMEM/F-12; ThermoFisher Scientific) supplemented with 10% fetal bovine serum (FBS; ThermoFisher Scientific) and 1% Antibiotic-Antimycotic (Anti-Anti; ThermoFisher Scientific) at 70°C under constant stirring at 500 rpm for 30 minutes to ensure complete dissolution. The LAP solution was then sterilized using a 0.22 µm filter pore. Sterile LAP and GelMA solutions were combined to obtain the desired final composition and then mixed at 40°C at 500 rpm for 15 minutes to ensure full mixing. GelMA/LAP solutions were stored at 4°C protected from light then pre-warmed to 37°C prior to use.

### Cell Culture and Seeding

Human airway epithelial cell lines (Calu-3) and human fetal lung fibroblasts (HFL1) were generously provided by Vancouver General Hospital (Vancouver, British Columbia) and St. Paul’s Hospital (Vancouver, British Columbia) respectively. Human umbilical vein endothelial cells (HUVECs) were purchased from American Type Culture Collection (ATCC, Manassas, VA, USA). Prior to experiments, each cell type was cultured in separate standard T75 cell culture flasks (Corning) containing the appropriate cell culture medium; DMEM/F-12 supplemented with 10% FBS and 1% Anti-Anti for Calu-3 and HFL1 and endothelial cell growth medium (EGM-2; Lonza) for HUVEC. All cells were maintained at 37°C and 5% CO_2,_ and the medium was changed every 2 days. Cells were harvested at approximately 80% confluency and prepared for experiments.

To create cell suspensions, Calu-3 (passage 15-21) or HUVECs (passage 5-10) were rinsed with PBS then detached from the flask surface using pre-warmed Trypsin/EDTA solution (0.05%; ThermoFisher Scientific). Cells were then pelleted by centrifugation at 1500 rpm for 5 minutes at room temperature, counted using an automated cell counter (Luna-II; Logos Biosystems) and resuspended in the appropriate cell culture media.

To create cell/gel mixtures, HFL1 (passage 4-8) were detached from the flask surface using pre-warmed Trypsin/EDTA solution (0.05%), pelleted by centrifugation at 1500 rpm for 5 minutes at room temperature, counted and resuspended in hydrogel pre-polymer solution.

### Hydrogel Selection and Optimization

#### Hydrogel Crosslinking Efficiency and Density

Disc-shaped hydrogel samples (15 mm diameter, 1.7 mm thickness) were used for sol fraction and swelling experiments. These experiments were conducted using established protocols^47,89^ and used to assess crosslinking efficiency and density. To create samples, 300 µL of a GelMA/LAP mixture was pipetted into polydimethylsiloxane (PDMS) molds and exposed to violet light (399 nm) (ColorBright, 365-460 nm, Flexfire LEDs, Cat. UV-24V-16FT) (∼8 mW/cm^2^) for 60 seconds. Directly after crosslinking, the hydrogels (6 per experimental group) were removed from the PDMS mold and weighed to obtain their initial wet mass (m_wet,t0_).

At this timepoint, 3 of the 6 hydrogels were transferred into individual containers, protected from light and frozen (-80°C) overnight. Frozen samples were then lyophilized in a freeze-dryer (Labcono) overnight to obtain the dry mass of the hydrogel (m_dry,t0_). The initial dry mass of the remaining 3 samples was then calculated using the actual macromer fraction and each sample’s starting wet mass. Splitting samples created at the same time point and gel thawing was intended to minimize sample variability. The actual macromer concentration [n = 3] and initial dry mass [n = 3] for samples that were not processed for measurement post-crosslinking were calculated based on the equation 1 and equation 2.

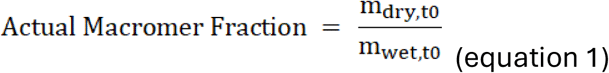

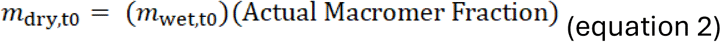

The remaining 3 samples that were not processed for sol fraction, were weighed immediately following crosslinking (m_wet,t0_) and then placed into wells of a 6-well plate. The samples were then submerged in 3 mL phosphate buffered saline (PBS) and incubated at 37°C for 24 hours to reach equilibrium swelling^90^. After 24 hours, the swollen hydrogel samples were blot dried with a KimWipe to remove the residual liquid, and their swollen mass measured (m_wet, t24_). Once measured, the samples were frozen, lyophilized and weighed to obtain the dry mass (m_dry, t24_). The swelling ratio for the hydrogels [n = 3] was calculated according to equation 3.

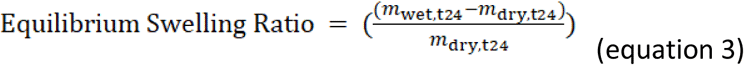

Sol fraction analysis was performed on all hydrogel compositions to analyze crosslinking capability. Samples measured for initial dry mass were also placed in PBS for 24 hours post weighing to remove unreacted polymer and determine sol fraction. When placed in an aqueous environment post-crosslinking, the polymer chains not attached to the network diffuse out of the sample resulting in a reduction in mass. This reduction in mass (sol fraction) reaches an equilibrium point after 24 hours when incubated at physiological conditions (37°C). After 24 hours incubation, PBS was quickly removed, and hydrogels underwent a freeze-drying cycle again. The final dry mass (m_dry,t24_) of samples was then recorded and sol fraction was calculated using equation 4.

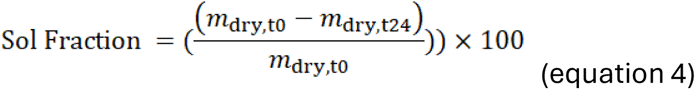

#### Cell-Laden Hydrogel Constructs to Assess Cell Encapsulation and Adhesions

Cell-laden hydrogel constructs were used to assess the cell encapsulation and adhesive characteristics of GelMA/LAP hydrogels. The wells of a standard 96-well plate (or wells of a custom 4-well glass bottom PDMS microdevice (**Fig. S4**) were used as a mold to create 3D disc-shaped constructs (6.5 mm diameter, 1.5 mm thickness for well plates and 4mm diameter, 0.8mm for 4-well PDMS-Glass microdevices) compatible with standard cell culture techniques. As shown in **Fig. S1**, hydrogel constructs were used to create various airway-related models to optimize the hydrogel formulation and ensure suitability for AOC integration.

Lung fibroblast (HFL1) laden hydrogel constructs (**Fig. S1a**) were created by mixing fibroblasts (1-5×10^6^ cells/mL) with the liquid pre-polymer GelMA solution. Cells were harvested, pelleted, and resuspended in GelMA prior to construct formation. The constructs were then formed by dispensing 50 µl of HFL1/GelMA mixtures into the wells of a 96-well plate and polymerized under violet light (399 nm) (8-10 mW/cm^2^) for 60 seconds. To generate constructs with varying mechanical and biochemical properties, nine different hydrogel formulations were prepared and analyzed. After polymerization, constructs were then topped with 100 µL of DMEM/F-12 cell culture medium supplemented with 10% FBS and cultured for 3-7 days in a standard culture incubator. Media was exchanged every 2 days. Morphology and viability of encapsulated cells was assessed.

Endothelial or epithelial monolayers were grown on the surface of cell-less bulk GelMA hydrogels as shown in the schematics in **Fig.S1b** and **Fig.S1c**. Cell-less constructs were created using the similar methods as fibroblast-laden constructs. However, fibroblasts were not mixed into the liquid pre-polymer GelMA solution prior to construct formation. Endothelial (HUVEC, 2 - 8×10^5^ cells/mL) or airway epithelial (Calu-3, 1 - 5×10^6^ cells/mL) cells were seeded on the hydrogel surface by adding 100 µL cell suspension (200µl for 4-well microdevices) to each well to coat the hydrogel. Media was exchanged every 2 days. Morphology and confluency of the cell monolayers were assessed.

Epithelial-fibroblast co-cultures (**Fig. S1d**) were created using similar protocols to monocultures. Lung fibroblasts (HFL1) were allowed to grow for 3 days before airway epithelial cells (Calu-3) were seeded on the surface of the fibroblast-laden hydrogel.

#### Viability: Fibroblasts encapsulated in hydrogel

The viability of fibroblasts encapsulated within hydrogel was assessed using dead cell and nuclei staining. Dead cell staining was performed using 4 µM Ethidium homodimer-1 (EthD-1) according to the Live/Dead Viability/Cytotoxicity Kit, for mammalian cells (Invitrogen). Hoechst 33342 at a 1:1000 dilution was used as a nuclear stain. The number of dead and total cells embedded in hydrogels made from different concentrations of GelMA were quantified using ImageJ from the EthD-1 and Hoechst-stained images respectively. Viability was calculated according to equation 5.

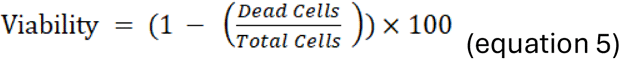

### Airway-On-A-Chip (AOC)

#### Microfluidic Device Design and Fabrication

The multilayered, multicompartment PDMS based microfluidic device builds upon the vertically stacked, bi-chamber design commonly used in organ-on-a-chip devices^24,26,28,32,33,38^. PDMS was selected for device construction as it supports real-time analysis via traditional microscopy as well as the opportunity to deconstruct the model and gain access to the hydrogel and membrane for off-chip histology or molecular analysis. The range of compatible assays and endpoints is expanded with an extractable gel, enhancing the functionality of the AOC and expanding the range of biological questions it can address.

The external microfluidic device consists of two PDMS layers containing rectangular microchannels (1 mm x 1 mm) separated by a thin (10 µm) porous (0.4 µm pores) polyester (PET) membrane. The microchannels contain a straight portion in the middle of the PDMS layer that creates the overlapping culture chambers but branches away on either side of the central microchannel to enable alignment with fluidic ports and ensure fluidic access to the chip. Fluidic ports were incorporated directly into the PDMS layers during the molding process using hollow needles (22G) as removable structures and the tips were aligned with the small straight portion of the branched section of the microchannel. **Fig. S5** visually demonstrates the creation of fluidic ports during the molding process. This approach ensures that the two chambers remain independently controllable when the device is assembled, facilitating precise control of flow (media or air) through the culture chambers and the establishment of a dynamic air-tissue interface.

In addition to the central microchannel, the bottom PDMS layer also contains lumen guide channels on either side of the central microchannel. The lumen guide channels were created during the molding processes in a similar manner to fluidic ports using hollow needles (25G) and connects the central portion of the bottom microchannel to the chip’s exterior. The guides operate as a mechanism to temporarily insert, fix, and extract an acupuncture needle enabling the microdevice to be compatible with needle based sacrificial molding techniques.

PET membranes were cut from commercially available Transwell® inserts. PDMS layers were generated through replica molding using 3D printed master molds. All molds were designed using SolidWorks and fabricated using SLA 3D printing (MiiCraft Ultra50, Creative CADWorks). Details on the master mold design and PDMS fabrication process are found in **S5** and illustrated in **Fig. S5**. To obtain an assembled AOC microdevice, the PDMS layers were exposed to oxygen plasma treatment (Harrick Plasma Cleaner) then manually aligned and bonded through thermal compression (24 hours at 60°C).

Fluidic connections consisting of 22G stainless-steel needles (McMaster-Carr) combined with Tygon® microfluidic tubing (Cole-Parmer) were inserted directly into the ports contained within the PDMS layers to couple the AOC microfluidic device with the flow management systems. To reinforce the fluidic connections for long-term cell culture, the fluidic connections inserted into the microchannels were encapsulated in a PDMS moat as described in^91^. Additional information on the moat design and fabrication process can be found in **S6** and visually shown in **Fig. S6.**

Prior to incorporating the ECM and cellular components, the assembled PDMS-based microfluidic device undergoes a three-step preparation process. The preparation process includes disinfecting the AOC by flushing the system with 70% ethanol followed by washing with sterile PBS to remove residual ethanol. Once disinfected, the PDMS-based AOC microfluidic device is equilibrated by loading the system with cell culture media and incubating it overnight in a humidified CO_2_ incubator.

#### Hydrogel Integration: Hydrogel Loading, In Situ Gelation and Lumen Patterning

The 3D hydrogel component of the AOC is incorporated once the external microfluidic device is assembled and prepared. A liquid hydrogel pre-polymer solution is manually loaded through the fluidic ports using a 1 mL syringe and once the bottom microchannel was completely filled, the hydrogel was polymerized *in situ* by exposing the hydrogel containing AOC microdevice to violet light (399 nm) (8 – 10 mW/cm^2^) for 60 seconds. To ensure uniform polymerization, the devices were exposed to light for 30 seconds on one side, then flipped and exposed for an additional 30 seconds. Post polymerization, the bottom microchannel loses its perfusion capabilities and no longer requires fluidic access so it is placed in a “closed loop” configuration by connecting the inlet and outlet tubing of the bottom microchannel to each other using a blocked 22G stainless steel needle.

A hollow circular channel (lumen) can be patterned directly into the hydrogel using an bovine serum albumin (BSA) coated acupuncture needle (0.25 mm gauge x 50 mm length; Dong Bang) as the template for sacrificial molding. Patterning is completed simultaneously with polymerization. The needle is inserted through the central portion of the bottom microchannel using the lumen guide channels. With the needle inserted, the liquid hydrogel pre-polymer solution can be introduced into the bottom microchannel, allowing it to fully surround the suspended needle, embedding the needle within the hydrogel matrix. After photopolymerization, the needle was gently withdrawn, leaving a hollow lumen structure within the hydrogel. Fluidic connections to the lumen were established by attaching Tygon® tubing to the 25G needles inserted into the lumen guide channel. Additional details on acupuncture needle-based sacrificial molding are provided in **S7** and the steps are depicted in **Fig. S7**.

### Flow Management Systems

To control the dynamic microenvironment within the AOC, the PDMS-based microfluidic device was interfaced with two independent, external pump-based flow management systems designed to mimic circulation and inhalation. The systems were used to precisely control microenvironmental conditions within each compartment and replicate physiological flows and shear stresses.

Although a recognized disadvantage of PDMS for organ-on-a-chip models is that it can absorb small molecules, a study conducted by Winkler and Herland^92^ highlighted that the material selected for the tubing connecting microfluidic devices to perfusion systems dominates over device sorption. In this work, care was taken to utilize appropriate tubing materials within the perfusion and airflow systems and minimize the overall tubing lengths to reduce absorption. In addition, since the particulates of interest associated with exposure studies are also significantly larger than small molecules, the limitations associated with small molecule absorption were deemed minimal for the device’s intended application. This is reflected in previous LOC models utilized for exposure studies being constructed from PDMS^29,30^.

#### Perfusion

To support the cell culture established within the AOC microdevice and mimic vascular perfusion including *in vivo* fluid shear stresses associated with blood flow, the AOC model was connected to a programmable, peristaltic pump-based perfusion system via fluidic connections integrated into the PDMS microfluidic device. Cell culture media was continuously drawn from inlet reservoirs via a peristaltic pump (Ismatec IPC ISM934C) and perfused through the device terminating at outlet reservoirs. The multichannel pump enabled up to 12 chips to be perfused simultaneously and was operated at a volumetric flow rate of 5 µL/min to mimic blood flow-induced shear stresses.

The AOC’s compartmentalized design facilitates continuous and independent sampling of effluent from the various compartments, permitting longitudinal analysis of cytokine secretion without disturbing the culture, an advantage highlighted by Benam et al.^27^. When operated under air-tissue interface conditions, effluent is collected from the vasculature compartment only, resulting in pooling of secreted factors (i.e. cytokines) limiting the ability to distinguish cytokine release origin to a specific cell-type.

#### Airflow

A vacuum pump-driven system was utilized to apply airflow to the upper microchannel to establish a dynamic air-tissue interface within the AOC. The airflow system (**Fig. S3.2**) consists of a HEPA filter, Tygon® tubing (¼” OD, McMaster Carr, Cat. 6516T17), two 5-way manifolds (McMaster Carr, Cat. 5779K318) with 22G needle connectors, and programmable vacuum pump (Gilian GilAir Plus, Cat. P/N 610-0901-02-R), arranged to support up to four AOC microdevices in parallel. This configuration enabled simultaneous exposure of multiple devices to physiological airflow conditions using a single pump. The vacuum pump was operated at 10 mL/min per chip, continuously drawing filtered air from the ambient environment through the upper microchannel and generating an estimated wall shear stress of approximately 0.2 dyn/cm^2^, comparable to that experienced by epithelial cells in the conducting zone of the airways^93,94^.

Environmental conditions, including humidity and temperature, were maintained by placing the perfusion system and AOC microfluidic devices inside a standard cell culture incubator during operation. This setup ensured that the cell culture medium remained at physiological temperature and pH throughout experiments.

### Microfluidic Cell Seeding of Airway-On-A-Chip

Microfluidic cell seeding was performed by manual syringe-based injection through fluidic connections inserted into the AOC microfluidic device. A cell suspension or liquid cell/gel mixture was loaded into a 1 mL syringe fitted with a 22G dispensing tip and connected to the open end of the Tygon® tubing based fluidic connections. The solution was slowly injected into the tubing until the channel was completely filled. The compartmentalized design enabled independent seeding of distinct cell types into separate compartments of the device. This enabled mono-, co-, and tri-cultures to be established within the AOC.

Prior to cell seeding, the separation membrane was coated with an ECM solution composed of collagen IV, laminin, and fibronectin by introducing the coating solution into the upper microchannel and incubating overnight at 37°C in 5% CO_2_.

After membrane coating, the stromal compartment was established within the bottom microchannel of the AOC. A HFL1/GelMA mixture (1-5×10^6^ cells/mL) was loaded into the bottom microchannel and polymerized *in situ* around a sacrificial acupuncture needle that was subsequently withdrawn, to create a hollow lumen within the hydrogel. The upper microchannel was then connected to the perfusion system, and the cells within the AOC were cultured under perfusion (300 µL/h) for 2 days.

After 2 days, the AOC microdevice was then disconnected from the perfusion system, and HUVECs (2-3×10^6^ cells/mL) were manually seeded into the lumen. HUVECs adhered to the interior surface of the lumen under static conditions, with the lumen fluidics closed off by connecting the inlet and outlet tubing using a blocked needle. Devices were incubated in a standard cell culture incubator at 37 °C in 5% CO₂. To facilitate adhesion within the 3D lumen, vascularized AOCs were rotated 90° hourly for 4 hours, following protocols used by previous groups^57,95^. Post-static incubation, the lumen and upper microchannel were connected to the perfusion system and cultured for 3 days at 300 µL/h with DMEM/F-12 and EGM-2, respectively, to establish a fibroblast–endothelial co-culture. After 3 days, the AOC microdevice was disconnected from the perfusion system, and the upper microchannel was seeded with Calu-3 cells (3–6 x10^6^ cells/mL). The inlet and outlet tubing of the upper microchannel were then connected using a blocked 22G needle, and the device was incubated in a standard cell culture incubator under static conditions for ∼4 hours to facilitate cell adhesion to the membrane surface. The AOC was then reconnected to the perfusion system and the cell-containing AOC microdevices were cultured under perfusion (300 µL/h) for 7 days in submerged state. Once the Calu-3 cells reached confluency, the media in the upper microchannel was aspirated to air-lift the Calu-3 cells and establish an air-tissue interface. The AOC devices were then cultured under a dynamic air-tissue interface (perfused at 300 µl/h) for an additional 12-15 days.

During the chip development process, monoculture and co-culture versions of the AOC were utilized. For these models, the cell seeding steps for cells not incorporated were skipped. The lumen patterning step was also skipped for submerged co-culture models as the upper channel was used for media supply and perfusion.

### Treatments and Exposures

To demonstrate the AOC system’s ability to recapitulate airway remodeling and be utilized for studying inhalation exposure, proof-of-concept studies were undertaken using TGF-B1 and whole wood smoke.

#### Transforming Growth Factor Beta 1 (TGF-β1) Treatment

To simulate upregulation of TGF-β1 in the airways associated with smoke exposure^68,69,73^ and validate the platform’s ability to respond to stimuli, AOC models were treated with TGF-β1 (50 ng/mL) for 72 hours. GelMA hydrogel constructs were also treated with TGF-β1 during the hydrogel optimization process to confirm that fibroblasts encapsulated within the selected GelMA/LAP formulation retained their ability to respond to stimuli.

TGF-β1 (STEMCELL Technologies) was reconstituted in 10 mM hydrochloric acid following the manufacturers’ protocol to create a 0.1 mg/mL solution, then diluted to 50 ng/mL in serum-reduced (1% FBS) DMEM/F-12 cell culture medium prior to use. Control samples were treated with an equivalent volume of fresh serum-reduced DMEM/F-12 cell culture medium without TGF-β1 for the same duration and conditions (static vs perfused) using the same formats (GelMA-based hydrogel construct vs AOC model). For GelMA hydrogel constructs, treatment was done under static conditions, while the AOC models were treated under perfusion with minimal shear (9.4×10^-4^ dyn/cm^2^).

For GelMA hydrogel constructs, once a healthy culture was established, the media covering the constructs was aspirated, and the constructs were washed manually with PBS (pre-warmed to 37°C) to remove traces of residual serum. Post-washing, the cells were serum-starved (DMEM/F-12 medium with 1% FBS) for 24 hours followed by TGF-β1 treatment for 72 hours.

TGF-β1 was added to the appropriate media reservoirs of the AOC model and coupled with the perfusion system. This allowed dynamic treatment of the cytokine via continuous perfusion through the top microchannel. The upper microchannel of AOC microdevices were washed with pre-warmed (37°C) PBS to remove residual serum, then serum-starved for 24 hours before TGF-β1 treatment for 72 hours. Serum starvation and treatment were conducted under continuous perfusion (300 µl/h and 60 µl/h, respectively), with a lower flow rate selected for the latter to reduce shear stresses and conserve reagents. Effluent samples were collected before and after treatment and stored at -80°C for downstream cytokine release and viability assays.

GelMA constructs and fibroblast laden AOCs (monoculture models), were fixed and stained for the myofibroblast marker α-smooth muscle actin (α-SMA) and filamentous actin (F-Actin) after treatment to visualize the cytoskeleton and observe changes in cell morphology and cellular behavior. Morphological and behavior changes were assessed by quantifying fibroblast activation and fibroblast-to-myofibroblast differentiation. Epithelial-fibroblast laden AOCs (co-culture models) were also treated with TGF-β1 and analyzed for cytotoxicity and cytokine release.

#### Whole Wood Smoke Exposure

To demonstrate the suitability of the AOC microdevice for exposure study applications, a protocol for direct on-chip exposure to whole wood smoke was established, and proof-of-concept studies were undertaken. Whole wood smoke was generated by combustion of cedar wood chips through a controlled system adapted from Kim *et al*.^96^ equipped with real-time monitoring capabilities including a particulate analyzer, and temperature and humidity sensors. Details on the wood smoke exposure setup are found in **S3. Fig. S3.1** highlights the components of the wood smoke generator, and **Fig. S3.2** demonstrates how the airflow system of the AOC is interfaced with the sampling port of the exposure chamber.

As shown in the schematic in **Fig. S3.2a**, smoke exposure on-chip was achieved by connecting the airflow system of the AOC to an exposure chamber for macroscopic human exposures coupled with a furnace-based wood smoke generator. When interfaced with the exposure booth, the vacuum pump-based airflow system draws whole wood smoke through the upper microchannel, exposing the apical surface of the epithelium that lines the microchannel in a biomimetic manner.

Prior to exposure, epithelial-lumen patterned interstitial fibroblast AOC models (co-culture with patterned lumen) were pre-treated with serum-reduced DMEM/F-12 cell culture medium for 24 hours before filling the top microchannel with air to establish an air-tissue interface. Wood smoke aerosol with a high PM2.5 concentration (480.68 ± 173.08 µg/mL) was generated by burning a set mass of dried cedar woodchip (4.25 ± 0.5 g) which was then continuously delivered to AOC models in a biomimetic manner for 1-2 hours. The exposure conditions were selected to reflect the ground-level PM2.5 levels that occur during forest fire events^97^ and are used in human *in vivo* wood smoke exposure studies ^72,98,99^. After exposure, AOC models were perfused for 24 hours before samples were collected and analyzed for cytotoxicity (LDH) and inflammatory markers (IL-6 and IL-8).

Environmental conditions, including humidity and temperature, were maintained by placing the airflow systems (excluding filters and pumps) and chips in a conventional CO_2_ incubator during exposures to ensure that wood smoke or ambient air reaching the cells contained within the AOC were maintained at 37°C. **Fig. S3.2c-d** illustrates the operational setup. Short tubing lengths and pre-conditioning of the tubes were used to reduce absorption and condensation, thereby reducing particulate loss during delivery.

### Microarchitecture Visualization

Fluorescent microbeads were used to visualize the 3D microarchitectures created within the AOC and illustrate the spatial arrangement achievable when cells are incorporated into the hydrogel bulk prior to polymerization. 1 µm red (542 nm) fluorescent polystyrene microbeads (ThermoFisher Scientific, Cat. R0100) were utilized to render the majority of the hydrogel fluorescent while 15 µm green (480 nm) fluorescent polystyrene microbeads (Bangs Laboratories Inc, Cat. FSDG009) were utilized to mimic fibroblasts embedded in the GelMA hydrogel. When 15 µm microbeads were used to mimic cells, the 1 µm microbeads were mixed with PBS and perfused through the lumen to visualize the hollow structure within the hydrogel and illustrate how HUVECs can be incorporated into the lumen interior.

### Immunocytochemistry Staining

Immunocytochemistry (ICC) staining was performed using standard techniques on fixed samples. Unless specified, all reagents used for ICC were purchased from ThermoFisher Scientific.

Prior to ICC staining, cell-laden constructs and AOC models were washed three times with PBS and fixed with 4% paraformaldehyde (PFA) for 1 hour at room temperature (RT). Post-fixation, residual fixative was removed by washing samples three times with PBS prior to blocking. Samples were blocked in 5% normal goat serum, 0.1% Triton X-100 in PBS for 1 hour at RT, then washed three times with PBS.

Primary antibodies for myofibroblast marker alpha smooth muscle actin (α-SMA, 1 µg/mL; Invitrogen, Cat. 14-9760-82) or vascular endothelial cadherin (VE-Cadherin, 5 µg/mL; Invitrogen, Cat. 36-1900) or Zona Occludens-1 (ZO-1, 5 µg/mL; Invitrogen Cat. 33-9100) or Mucin 5AC (MUC5AC, 2 µg/mL; Invitrogen Cat. MA5-12178) were incubated with samples overnight at 4°C.

After overnight incubation, samples were washed three times with PBS, then stained with secondary antibody (2 µg/mL, Goat Anti-Mouse Alexa Fluor 555 (Invitrogen Cat. A28180) or 4 µg/mL Goat Anti-Rabbit Alexa Fluor 555 (Invitrogen Cat. A-21428) for 1 hour at RT. Post-ICC staining, samples were counterstained with Alexa Fluor 488-conjugated phalloidin and nuclear stain (DAPI or Hoechst 33342) to visualize the cytoskeleton and visually assess morphology. For AOC models, washing and staining were performed on-chip by manually perfusing reagents through the microchannels. Reagents were loaded into the top microchannel of AOC devices via fluidic connections and clamped during incubation. To prevent evaporation, samples were placed in a humidity chamber throughout the staining process. To enhance visualization, hydrogel samples were extracted from well plates (hydrogel constructs) or AOC devices post-staining and then mounted onto a glass coverslip. For hydrogel samples in the 4-well microdevices, extraction was not necessary as the samples were already in direct contact with the device’s glass coverslip bottom.

### Microscopy

Images were captured using either fluorescence (Nikon Eclipse TE2000-U Inverted Microscope) or confocal laser scanning microscopy (Zeiss Microscope) using 10×, 20× or 40× objective lens. To reduce variation, samples from each independent experiment were imaged in a single session. For confocal images, the bottom and tops of gel were identified as the slices where the number of nuclei was very low. Maximum intensity projection (MIP) confocal images were created in ImageJ.

Mean fluorescence intensity was quantified using ImageJ, while fibroblast morphology was assessed using a custom Python script that utilizes eccentricity to classify cells as elongated or spherical. Elongation was quantified as the percentage of cells exhibiting an elongated morphology. Since fibroblast nuclei are not perfectly spherical, an eccentricity threshold of ∼ 0.5 was chosen to classify nuclei as spherical.

### Cytokine Release

The levels of IL-6 and IL-8 in effluent samples collected from AOC models pre-and post-treatment or exposure were quantified via ELISA (IL-6 & IL-8; R&D Biosystems, Cat. DY206 and DY208) following the manufacturer’s protocols. Optical density measurements were obtained using SpectraMax i3 Plate reader and a four-parameter logistic curve based on standard curves was used to convert absorbance values into concentrations (pg/mL) using GraphPad Prism 9 Software. The standard curve for IL-6 ranged from 9.38 pg/mL to 600 pg/mL, while the standard curve for IL-8 ranged from 31.3 pg/mL to 2000 pg/mL. Samples were diluted by ½ (IL-6) or ¼ (IL-8) prior to assaying to fall within the detection range of the standard curve and were then multiplied by the dilution factor after reading to obtain cytokine concentration.

### Cytotoxicity

Cytotoxicity of AOC cultures was quantified by measuring the level of lactate dehydrogenase (LDH) levels in effluent samples collected pre- and post-treatment or exposure, using the CyQUANT LDH Cytotoxicity Assay (Invitrogen, Cat. 20300). LDH is released by cells into the extracellular space when they are damaged or injured, which is comparable to LDH release into culture media in *in vitro* models, thereby providing a means to indirectly monitor cytotoxicity over time without compromising the cell culture. Following the manufacturer’s protocol, 50 µL of collected effluent samples were transferred into a 96-well plate. The reaction mixture was added and the contents gently mixed by pipetting. After a 30-minute incubation, the stop solution was added, and absorbance was measured at 490 nm and 680 nm using a SpectraMax i3 Microplate Reader. LDH release was calculated by subtracting the 680 nm background absorbance from the 490 nm signal. To assess the cytotoxic effects of treatment/exposure, the LDH levels in effluent samples (determined as corrected LDH absorbance) collected from TGF-β1 treated or wood smoke-exposed chips were compared to controls.

### Statistical Analysis

For hydrogel optimization, differences in the means of sol fractions, swelling ratios, and viability were detected by a one-way ANOVA and subsequent Bonferroni post-hoc test using GraphPad Prism 9 Software. Differences were considered significant when p < 0.05.

For treatment and exposures studies, differences between means of control and experimental (TGF-β1 treated or whole wood smoke exposed) were determined by two tailed t-tests using GraphPad Prism 9 Software, with a treatment/exposure being considered significant when p < 0.05. F-tests were performed to determine whether a student’s or Welch’s t-test was performed.

## Acknowledgements

The authors would like to acknowledge Dr. Don Sin and Dr. Tillie Hackett for supplying cells, Jung Yi Cau for assisting with PDMS based fabrication and Karoline Moo for assisting with setting up the initial wood smoke system. They would also like to acknowledge UBC’s Faculty of Medicine funding provided through the Multidisciplinary Research Program in Medicine and Dr. Min Hyung Ryu, Bora Jin, and Hana Kim. As well as funding provided by Mitacs IT13602, Providence Health Care, the Natural Sciences and Engineering Research Council of Canada, and the Canadian Institutes of Health Research.

## Author Contributions

Conceptualization, Tanya Bennet and Avineet Randhawa; Data curation, Tanya Bennet; Formal analysis, Tanya Bennet; Funding acquisition, Christopher Carlsten and Karen Cheung; Investigation, Tanya Bennet, Avineet Randhawa, Ryan Huff, Carley Schwartz and Yu Xu; Methodology, Tanya Bennet, Avineet Randhawa, Eric Lyall, Ryan Huff and Carley Schwartz; Software, Eric Lyall; Supervision, Christopher Carlsten and Karen Cheung; Validation, Tanya Bennet; Visualization, Tanya Bennet; Writing – original draft, Tanya Bennet; Writing – review & editing, Tanya Bennet, Tara Caffrey, Tanya Solomon and Karen Cheung.

## Data Availability Statement

The datasets used/or analysed during the current study available from the corresponding author on reasonable request.

The data analysis script used for elongation is available in the repository on GitHub that can be accessed through https://github.com/ericlyall/BioMEMS-Cell-Morphology-v0

