## Supplementary Information for "Engineering a Multilayer Microfluidic Airway-On-A-Chip with Tunable GelMA Hydrogel for Physiologically Relevant Aerosol Exposure Studies"

### S1. Cell Laden Hydrogel Construct Configurations

As shown in **Fig. S1**, well-based GelMA hydrogel constructs were used to create various *in vitro* airway models including a fibroblast-laden hydrogel, endothelial monolayer on a hydrogel surface, epithelial monolayer on a hydrogel surface, and co-culture containing an epithelial monolayer on the surface of a fibroblast-laden hydrogel. These simplified models were utilized for cell-based experiments conducted during the hydrogel optimization process to assess the suitability of GelMA/LAP formulations for AOC integration.

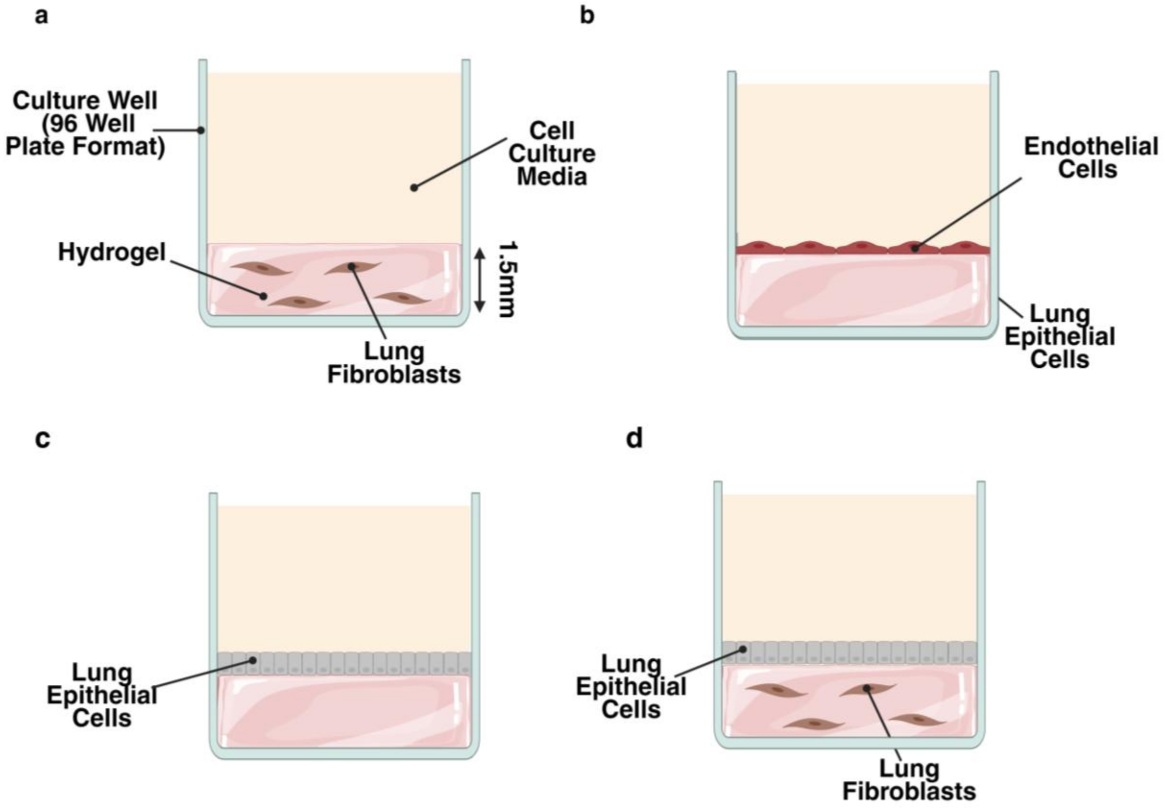

***Fig.S1.*** *Cell-laden hydrogel constructs (a) Fibroblasts encapsulated in hydrogel. Fibroblasts are exposed to all elements of the photopolymerization process as cells are mixed into the hydrogel that is photopolymerized in the well. (b) Endothelial monolayer on cell-less hydrogel surface. Hydrogel is photopolymerized prior to the cells being introduced. (c) Epithelial monolayer on cell-less hydrogel surface. Hydrogel is photopolymerized prior to the cells being introduced. (d) Epithelial-Fibroblast co-culture with fibroblasts encapsulated in the hydrogel and an epithelial monolayer on the surface of the hydrogel. Fibroblasts are exposed to photopolymerization process as cells are in hydrogel prior to polymerization but epithelial cells are introduced post-polymerization. Created with BioRender.com*

#### S2. Co-Culturing Capacity

The co-culturing capacity of the hydrogel is essential for enhancing physiological relevance and studying cell-cell and cell-ECM interactions. Although a simplified model, the epithelial-fibroblast co-culture hydrogel constructs (**Fig. S1d**) serve to further demonstrate the suitability of the 6% GelMA/0.6% LAP hydrogel for integration into the AOC platform, as well as provide preliminary evidence for potential future advancement of the AOC based on leveraging the hydrogel as the sole cell culture surface.

The confocal images and 3D reconstruction shown in **Fig. S2a** highlight how the GelMA hydrogel supports a 3D culture with direct cell-ECM contact, that mimics the epithelial-stromal interface of the small conducting airways. The 2D epithelial monolayer (**Fig. S2ai**) that forms on the hydrogel surface displays cobblestone morphology and appears to be tightly packed together, characteristics that are similar to the epithelium that lines the luminal surface of the airway. The fibroblasts encapsulated within the 3D hydrogel **(Fig. S2aii**) display characteristic elongated morphology and mimic resident fibroblasts found within the interstitial ECM. Unlike synthetic membranes that require additional ECM coatings, the GelMA hydrogel inherently supports cell attachment due to the retained cell-responsive motifs allowing the epithelial cell layer to be anchored directly to the underlying fibroblast-laden ECM hydrogel as illustrated in **Fig. S2aiii**.

Compared to fibroblast-laden hydrogel monocultures, fibroblasts co-cultured with epithelial cells within the GelMA hydrogel displayed enhanced elongation, increasing from 83.28 ± 1.15% to 89.08 ± 3.27% of fibroblasts displaying elongated morphology (**Fig. S2b-c**). These results highlight the role of epithelial-fibroblast communication and the importance of capturing cell-cell interactions within the AOC.

Similar to the fibroblast-laden hydrogel monocultures, the epithelial-fibroblast co-culture was treated with TGF-β1 to demonstrate the hydrogels suitability for AOC integration and its potential for studying cell-ECM and cell-cell interactions associated with airway remodeling, fibrosis and smoke exposure. Fibroblasts co-cultured with airway epithelial cells treated with TGF-β1 were also able to undergo fibroblast-to-myofibroblast differentiation, supported by morphological changes and an increase in α-SMA expression (**Fig. S2d-f**).

As shown in **Fig. S2g,** epithelial cells within the co-cultures also exhibited TGF-β1 induced morphological changes and barrier dysfunction. Epithelial cells lost typical cobblestone morphology and decreased in confluency (**Fig. S2h**) due to the formation of gaps within the monolayer indicating disruption of cell-cell adhesions and may be a preliminary sign of epithelial-to-mesenchymal transition.

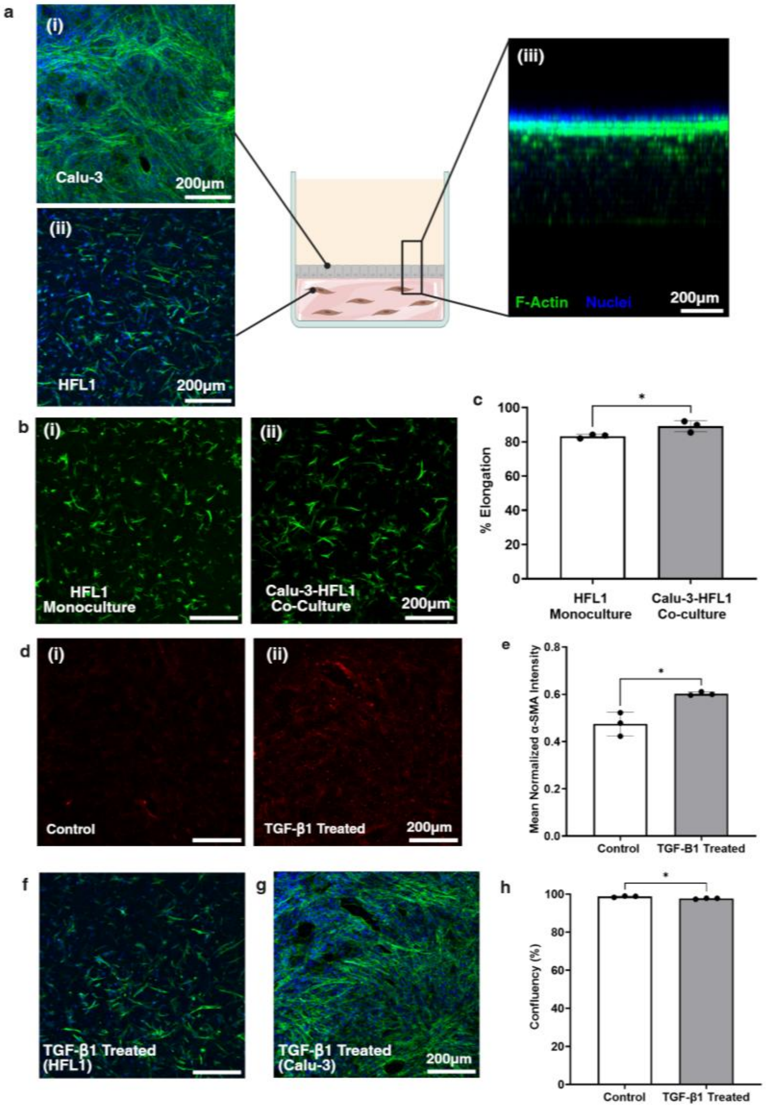

***Fig. S2.*** *Airway epithelial-lung fibroblast co-culture GelMA hydrogel construct. (a) Representative confocal images of the layers of the airway epithelial cells (Calu-3)-lung fibroblasts (HFL1) laden hydrogel co-culture. Cells were cultured on and encapsulated within a hydrogel composed of 6% GelMA/0.6% LAP. Cells were stained with Alexa Fluor 488-conjugated phalloidin (green) and Hoechst 33342 (blue). Schematic created with BioRender.com. (i) Calu-3 cells form a confluent monolayer with cobblestone morphology on the hydrogel surface. (ii) HFL1 cells encapsulated within hydrogel show elongated cell morphology. (iii) Side view of 3D confocal stack reconstruction highlighting 2D epithelial monolayer on hydrogel construct surface with fibroblasts dispersed throughout the 3D hydrogel. (b) Morphology of HFL1 cells embedded within 6% GelMA/0.6% LAP hydrogel when cultured (i) alone or in (ii) co-culture with Calu-3 cells. For co-culture, MIP excludes slices containing Calu-3 cells to improve visualization of fibroblast morphology. (c) Quantification of percentage of fibroblasts displaying elongated morphology (elongation) in GelMA hydrogel constructs. (d-g) TGF-β1 Treatment. Co-cultures treated with DMEM/F-12 plus 10% FBS without (control) or with TGF-β1 (50 ng/ml) for 72 hours. (d) Representative MIP confocal images of fibroblasts encapsulated in 6% GelMA/ 0.6 % LAP constructs stained for α-SMA (red). (e) Quantification of mean fluorescence intensity of α-SMA positive cells normalized to total cells. An increase in intensity indicates fibroblast differentiation into myofibroblasts. (f-g) Representative MIP confocal images of TGF-β1epithelail-fibroblast co-cultures. Representative MIP confocal images of control samples used for analysis are found in a. (f) Representative MIP confocal image of fibroblasts encapsulated through GelMA hydrogel highlighting morphological change towards star or web- morphology. Slices containing Calu-3 monolayer are eliminated from MIP. (g) Representative MIP confocal images of TGF-β1 treated Calu-3 monolayer highlighting morphological changes and reduction in monolayer confluency due to gap formation. Slices containing fibroblasts are eliminated from MIP. (h) Quantification of Calu-3 monolayer confluency. For each sample, a z-stack (25 μm step size) at three regions of interest was acquired using a 10x objective. Data represents mean ± standard deviation (n=3).* P values calculated using two tailed t-test. * *$p < 0.05$ and **$p<0.01$.*

### S3. Woodsmoke Exposure Setup

The woodsmoke exposure setup consists of a wood smoke generator, exposure chamber, and airflow system interfacing with the AOC platform.

#### 3.1 Wood Smoke Generator

As shown in **Fig. S3.1**, smoke generation was achieved using a quartz-tube furnace system consisting of a ceramic heating element mounted on a movable track that progressively moves along the length of a quartz tube to burn woodchips and generate constant, stable whole wood smoke conditions. To assist in uniform combustion and maintain PM2.5 levels within a desired range, dried cedar woodchips (Home Depot, Cat. 1000729924) were cut to a uniform size and evenly distributed inside the tube along its length. The linear actuator that controls the movement of the heating element was also set to a slow speed (1 cm/min) to maintain smoke generation for the entire exposure duration.

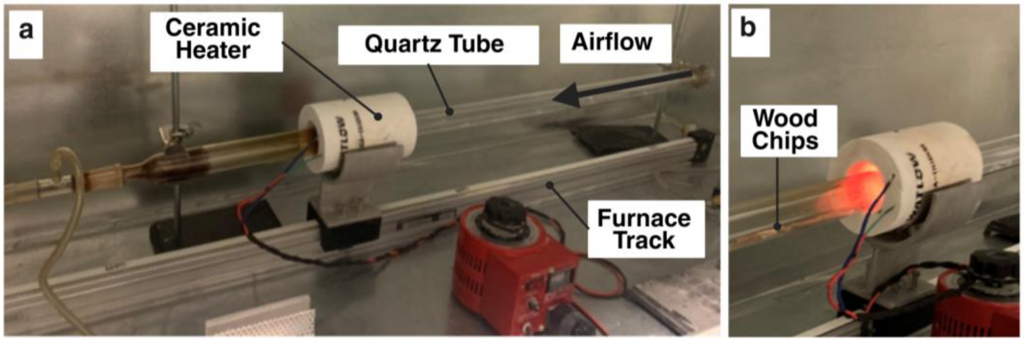

**Fig. S3.1.** Wood smoke generator (a) The smoke generator consists of a ceramic heater that surrounds a quartz tube and is mounted on a movable track. (b) To generate smoke, cedar wood chips are placed within the quartz tube and burnt as the heater progressively moves along the furnace track.

#### 3.2 Interfacing Airway-On-A-Chip with Exposure Chamber

Generated smoke was passed from the wood smoke generator to a human exposure chamber depicted in **Fig. S3.2a-b** via a secondary dilution system to obtain the desired concentration. Real-time, continuous monitoring of the environmental conditions (temperature and relative humidity) and smoke properties (PM2.5 and gas phase constituents (i.e. CO, CO2) was accomplished via particulate analyzers and sensors coupled to the sampling ports integrated into the chamber.

For exposure experiments, two configurations of the airflow system were utilized; one for filtered air and one for aerosolized wood smoke. To interface the AOC platform with the exposure chamber, the inlet tubing connected to the 5-way manifold of the airflow system was connected to a single port on the sampling manifold of the exposure chamber. In the filtered air (control) configuration, the inlet tubing was left uncoupled from the exposure chamber, and a HEPA filter was placed on the system opening to minimize contamination risks. The vacuum pump located downstream of the AOC controls delivery by drawing a stream of aerosolized wood smoke from the exposure chamber or filtered ambient air from the environment through the AOC microchannel without disrupting the air-tissue interface.

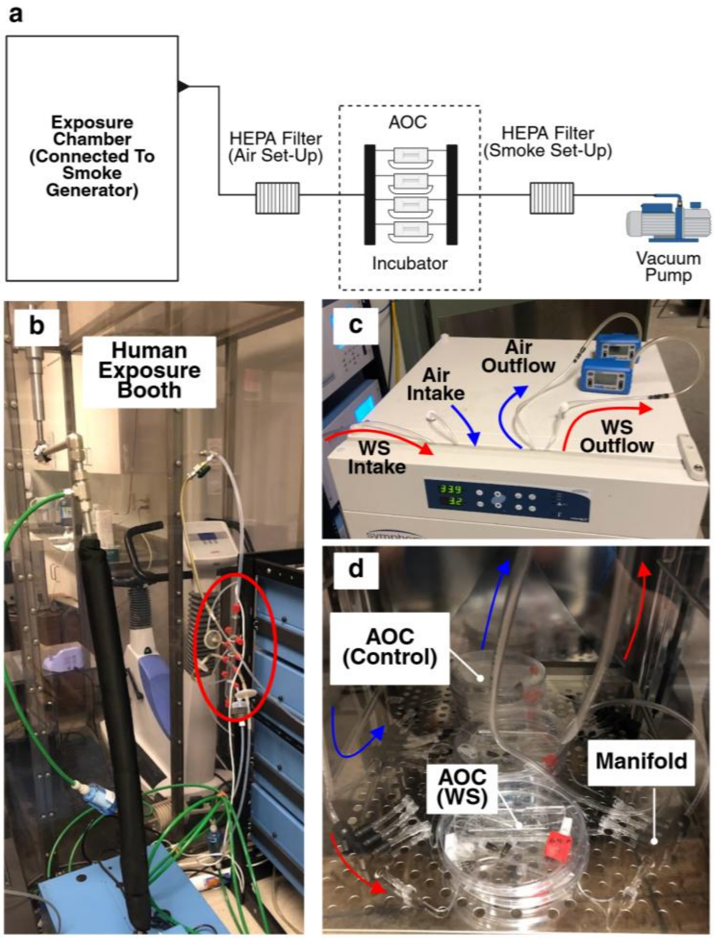

***Fig. S3.2.*** *Operational setup for wood smoke exposures (a) Schematic of the wood smoke exposure system. The airflow system is connected to the wood smoke exposure chamber via a sampling manifold. The exposure chamber is located downstream from the furnace-based wood smoke generator and upstream of the airflow system. HEPA filters were integrated in line with the AOC either upstream (for control configuration) or downstream (for exposure configuration) to eliminate contamination or protect the pump respectively. (b) The sampling port (highlighted with a red circle) on the exposure chamber is used to couple the airflow system of the AOC to the wood smoke generator. (c-d) During operation the (c) vacuum pumps and tubing connections to the air/smoke sources are left outside an incubator while (d) the remaining components of the airflow and exposure systems are placed into a standard cell culture incubator to control environmental conditions.*

### S4. 4-Well PDMS-Glass Microdevice

Polydimethylsiloxane (PDMS)-glass microdevices (**Fig. S4**) were created to enhance the microscopy compatibility of hydrogels and create hydrogel samples of decreased thickness. The microdevice consists of a PDMS gasket patterned with 4 independent gel wells (4mm diameter, 0.8mm height) overlaid with independent media reservoirs bonded to a glass coverslip. The total surface area of the cell culture well (gel well and media reservoir) is double that of a standard 96 well plate well, however the gel surface represents only 1/5th of the total surface area. To create the PDMS gaskets liquid PDMS (Sylgard 184, Dow Corning) mixed at a 10:1 (elastomer base: curing agent) ratio was poured into molds created using 3D printing (MiiCraft). A transparency film lid was placed over the PDMS-filled molds after degassing prior to curing to obtain a uniform thickness as previously described in[1].

To create gel samples within the microdevice liquid GelMA (10μl) can be loaded independently into each well and crosslinked *in situ*. This enables the hydrogel to be in direct contact with the glass coverslip. The media reservoir above the hydrogel can be filled with a cell suspension (for seeding) or DMEM/F-12 (for culture maintenance) to submerge the hydrogel and support cell seeding and culture.

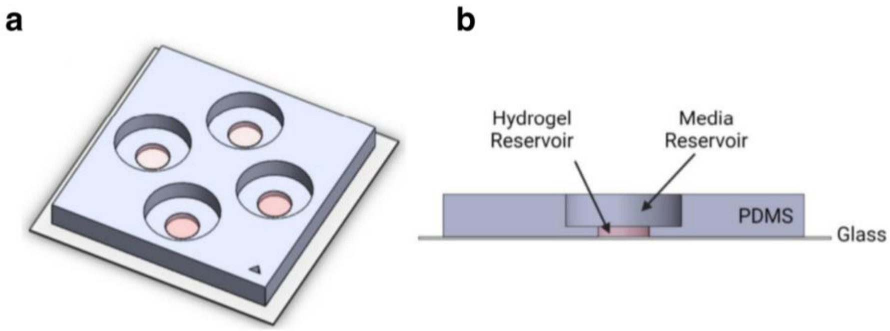

**Fig. S4.** 4 well PDMS-glass microdevice. (a) CAD rendering of assembled microdevice. (b) Cross sectional view of microdevice highlighting hydrogel reservoir overlaid by the larger media reservoir

#### S5. Airway-On-A-Chip Master Mold Design

The 3D printed master mold was design as the inverse of the resulting PDMS layers and contained through holes (0.56mm diameter) on the side walls allowing needles to be inserted into the molds and casted allowing fluidic ports (inlet/outlet) to be incorporated directly into the PDMS layer. This approach eliminates the need to create inlets/outlets via biopsy punches post fabrication, therefore minimizing the chance of misalignment of ports and channels and facilitating compatibility with microscopy as the in-plane connections do not impede microscope observation as they sit perpendicular to the imaging surface. Molds for the top PDMS layer contains one set of holes on either side of the upper microchannel extruded feature while the mold for the bottom PDMS layer contains two sets of holes, one on either side of the bottom microchannel extruded features and one on either side of the central portion of the microchannel extruded feature. To create fluidic connections for the microchannels, 25G stainless steel needles (McMaster-Carr, Cat. 75165A687) were inserted through the holes in the master mold and aligned with the extruded features corresponding to the microchannels. To create the lumen guide channels, two 25G needles were inserted and aligned with the central portion of the extruded feature. Once needles were inserted into the master mold, liquid PDMS (Sylgard 184; Dow Corning) mixed at a 10:1 (elastomer base: curing agent) ratio was poured into the molds to cast the PDMS layers. The PDMS filled molds were degassed in a desiccator for 30 – 40 min to remove bubbles and to obtain uniform thickness, a transparency film lid was used as previously described in[1]. PDMS layers were cured at 65° C overnight then carefully removed from the molds by first peeling the transparency film to reveal a flat PDMS layer and then separating the layers from the molds using a scapula. A schematic illustrating the microfabrication process can be seen in **Fig. S5**.

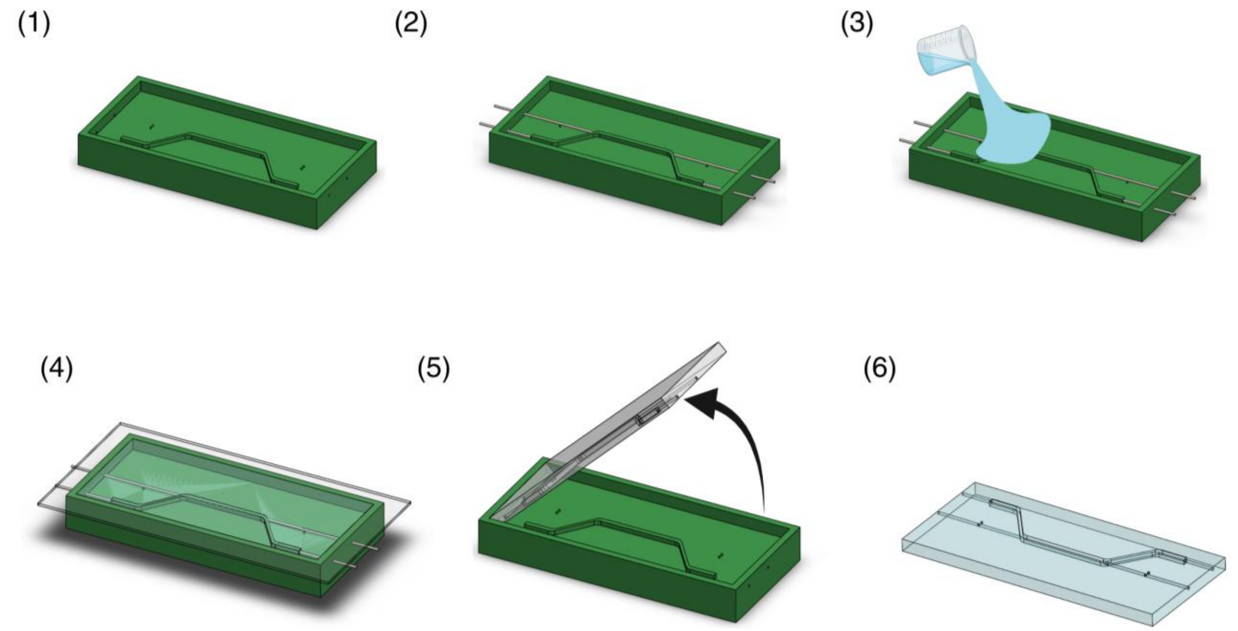

**Fig S5.** Fabrication of PDMS layers from 3D-printed master molds. (1) The 3D printed master molds contain holes within the side walls to enable needle insertion. (2) Needles are inserted into the master mold and aligned with the microchannel inlet/outlet structure or the central portion of the extruded microchannel. (3) Liquid PDMS is poured into the master mold until it slightly overflows. (4) After degassing, a transparency film is placed over the mold and then cured. (5) PDMS layers are removed from the mold. (6) Once removed from the molds, the PDMS layers are ready for assembly. Created with BioRender.com.

### S6. PDMS Moat Design

To support the long-term culture timelines required to establish differentiated airway epithelium, a PDMS moat system[1] that encapsulates the AOC device and fluidic interconnections was used to reinforce and increase the systems overall integrity. When operated without a patterned lumen such as in submerged co-cultures a simplified moat design utilizing a rectangular petri dish (**Fig. S6a**) as a mold was used. For models containing lumens a custom mold is used to secure the AOC device and tubing in place (**Fig. S6b**). 25G needles were inserted through holes in the custom mold and aligned with lumen channels of AOC devices. The needles were then pushed into the lumen channel to prevent seepage and liquid PDMS poured into the mold embedding the AOC microfluidic device within the PDMS moat. Once cured, the needles were removed and AOC devices with PDMS moat reinforcement were removed from mold. Fluidic interconnections attached to the top and bottom microchannels are encapsulated into PDMS moat and are able to be immediately connected to the flow management systems, while access to lumen channel remains accessible and compatible with lumen patterning.

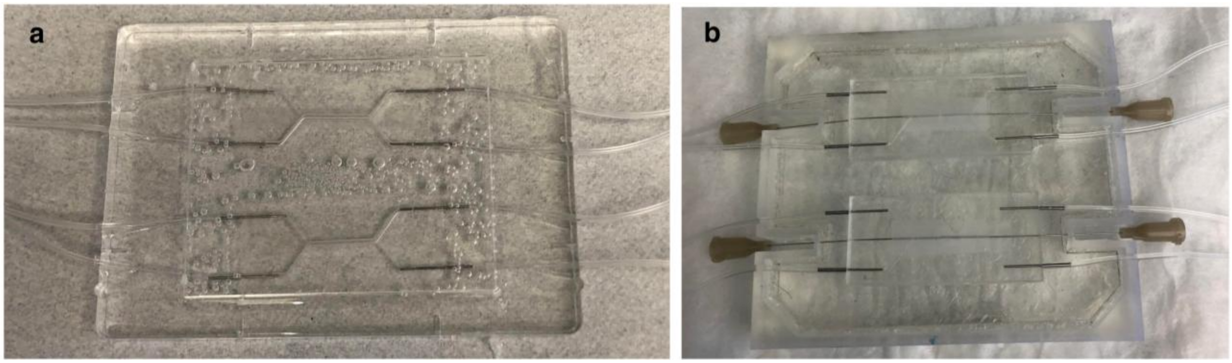

***Fig. S6.*** *PDMS moat reinforcement for long term cell culture (a) Co-culture without patterned lumen format. Microdevices are reinforced by placing assembled devices with fluidics into a petri dish before encapsulating connections in PDMS moat. (b) Lumen patterned and tri-culture format. Microdevices which contain additional ports for lumen patterning are reinforced by placing assembled devices into custom 3D printed mold. 25G needles are inserted through holes on the side walls of the mold and into lumen guide ports prior to casting the PDMS to ensure lumen guide channels remain open and accessible. For both formats, the models are reinforced in pairs to conserve PDMS and increase ease of handling.*

#### S7. Acupuncture Needle Based Sacrificial Molding

Acupuncture needle-based sacrificial molding utilizes a needle as a removable template. To create a hollow cylindrical structure directly within the GelMA hydrogel, the photopolymerizable hydrogel is crosslinked around the needle permanently capturing the needle geometry within the hydrogel. Removing the needle post crosslinking reveals a hollow structure. To facilitate lumen patterning, lumen guide channels were placed on either side of the central channel in the bottom PDMS layer. The guide channels allow hollow rigid 25G needles (12.7mm in length) to be permanently inserted into the PDMS layers, providing physical support and fluidic access to the central portion of the bottom microchannel. Prior to hydrogel loading, 25G needles are inserted into the two lumen guides channels on either side of the central portion of the bottom microchannel so that one end of the needle remains external to the microdevice, while the other enters the central portion of the bottom microchannel. These needles operate as a support structure for acupuncture needle insertion and fluidic connections. Once inserted, an acupuncture needle can be passed completely through the entire length of the central microchannel and temporarily fixed in a desired position within the bottom microchannel.  Prior to inserting the acupuncture needle, the needles were soaked in 1% BSA solution for 1 hr to prevent undesired adhesion between the hydrogel and the needle surface minimizing the potential for gel tearing during needle extraction and defects in the lumen structure. Once the acupuncture needle was fixed within the microdevice, the liquid GelMA pre-polymer solution was loaded into the bottom microchannel via manual perfusion. The hydrogel solution completely fills the bottom microchannel and freely flows around the needle, completely embedding the acupuncture needle within the bulk hydrogel structure. The hydrogel can then be photopolymerized by exposing the chip to violet light (399nm) for 60 s. Post polymerization, the needle is slowly withdrawn from the device using sterile tweezers revealing a hollow lumen directly within the 3D hydrogel. To stabilize the PDMS device during extraction, a second set of tweezers is placed on the device. Once formed Tygon® tubing can be attached to the 25G needles inserted into the guides to interface the lumen to the external pump and control fluid flow through the lumen. The step-by-step process for sacrificial molding using the AOC microdevice is shown in **Fig. S7**.

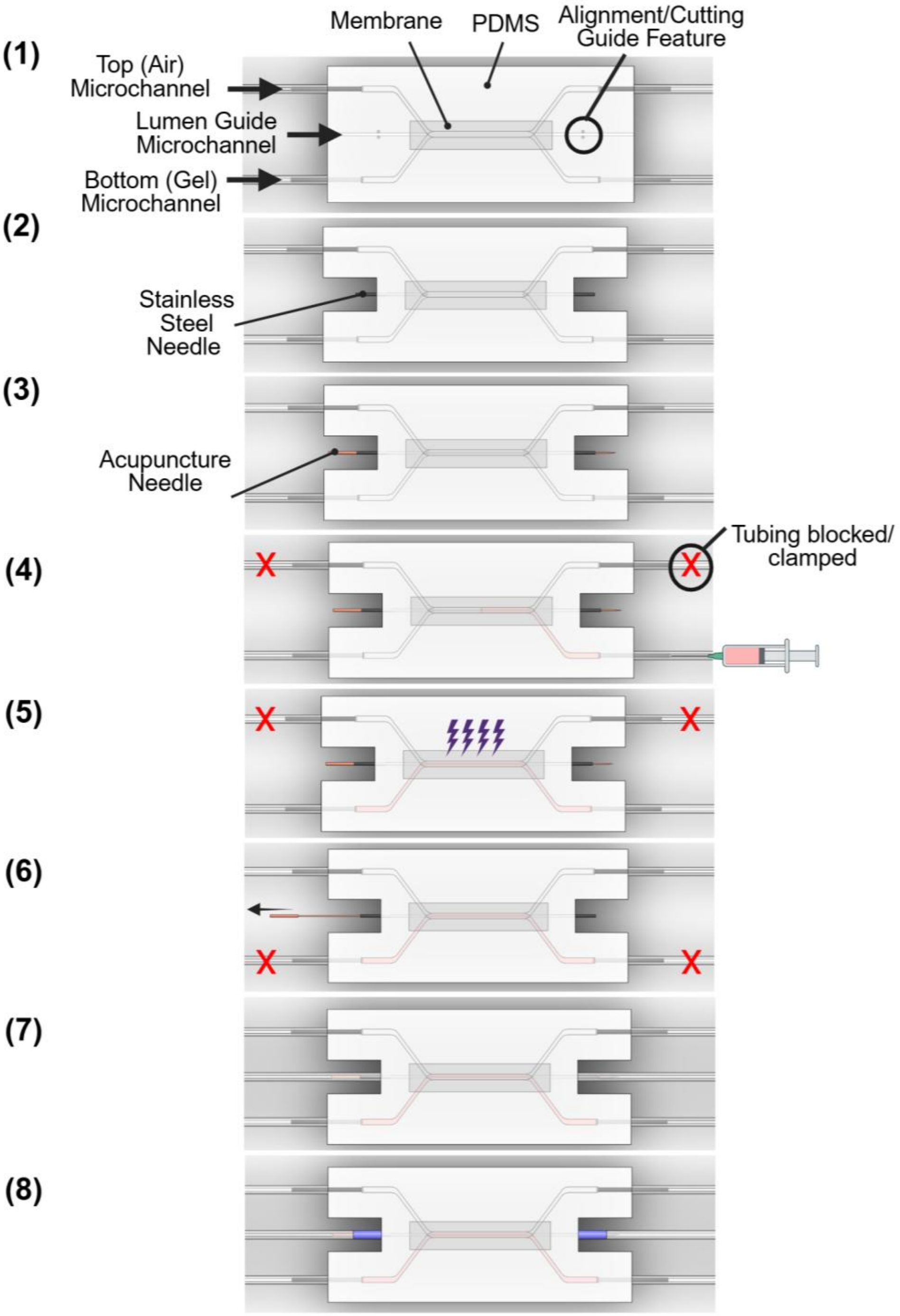

***Fig. S7.*** *Needle-based sacrificial molding process to create lumens in airway-on-a-chip microdevices. (1) After assembly, the AOC microfluidic device is ready for patterning. (2) PDMS was removed around lumen guide channels and needles inserted into the inlet and outlet of lumen guide channels until they enter the bottom (gel) microchannel. (3) The sacrificial needle was inserted into the microfluidic device through lumen guide channels. The needle is inserted on one side until it protrudes out the opposite side. (4) Fibroblast laden GelMA pre-polymer solution loaded into the bottom (gel) microchannel. (5) The AOC device was crosslinked by exposure to light. (6) Sacrificial needle is removed from the AOC device revealing a hollow lumen embedded within the fibroblast-laden GelMA hydrogel. (7) Fluidic tubing was connected to inlet/outlet needles to connect the lumen to the perfusion system. (8) Heat shrink tubing was used to reinforce inlet/outlet tubing connections to prevent leakage. Created with BioRender.com.*
